# Can a flower color ancestral polymorphism transcend speciation?

**DOI:** 10.1101/2023.11.14.566988

**Authors:** Mercedes Sánchez-Cabrera, Eduardo Narbona, Montserrat Arista, Pedro L. Ortiz, Francisco J. Jiménez-López, Amelia Fuller, Benjamin Carter, Justen B. Whittall

## Abstract

Polymorphisms are common in nature, but they are rarely shared among closely related species. They could originate through convergence, ancestral polymorphism, or introgression. Although shared neutral genomic variation is commonplace, very few examples of shared functional traits exist. The blue-orange petal color polymorphisms in two closely related species, *Lysimachia monelli* and *L. arvensis* were investigated with UV-vis reflectance spectra, flavonoid biochemistry, and transcriptome comparisons followed by climate niche analysis. The similarities in reflectance spectra, biochemistry, and transcriptomes suggest a single shift from blue-to-orange shared by both lineages is possible. Transcriptome comparisons reveal two orange-specific genes are directly involved in both blue-orange color polymorphisms: *DFR-2* specificity redirects flux from the malvidin to the pelargonidin while *BZ1-2* stabilizes the pelargonidin with glucose, producing the orange pelargonidin 3-glucoside. The climate niches for each color morph are the same between the two species for three temperature characteristics but differ for four precipitation variables. We suggest that this persistent flower color polymorphism may represent an ancestrally polymorphic trait that has transcended speciation with some unique ecological effects.

## Introduction

Since the Modern Synthesis, we have assumed that most phenotypic variation emerges within populations and is subsequently acted upon by evolutionary forces within species (Dobzhansky, 1937; Grant, 1981). Herein, we describe an ancestral flower color polymorphism that may have transcended the species boundary providing a concrete example of unsorted ancient phenotypic variation in two species with contrasting ecological effects. This study addresses how shared traits evolve across lineages, a central topic in understanding the evolution of biological diversity.

Polymorphisms, or variation among individuals of the same species, provide rare windows into the process of adaptation and speciation (Allison, 1961; Igić et al., 2008; Martin et al., 2013; Comeault et al., 2015). Polymorphisms among individuals in a population or among populations of a single species, are omnipresent across the tree of life (McKinnon & Pierotti, 2010; McLean & Stuart-Fox, 2014) and can be maintained by a diversity of forces including negative frequency-dependent selection (Madsen et al., 2022), heterozygote advantage (Sirugo et al., 2014), genetic drift (Wright, 1943, but see Schemske & Bierzychudek, 2007), gene flow (Martin et al., 2013) and spatially or temporally variable selection (Svensson, 2017). Some polymorphisms are phylogenetically dispersed (e.g. bird plumage color or flower color; Hugall & Stuart-Fox, 2012), while in others reoccur in very closely related species (e.g. heterostyly in *Primula*, shell chirality in *Amphidromus*, cryptic body color in *Timema*, and wing patterning in *Heliconius*; reviewed in Jamie & Meier, 2020). The molecular underpinnings, persistence mechanics and evolutionary forces acting on these rare cases of shared variation are largely unexplored.

Some studies have proposed that polymorphisms promote the utilization of a wide range of environmental resources and can act as a precursor to speciation which then become fixed after divergence (Corl et al., 2010; Hugall & Stuart-Fox, 2012; McLean & Stuart-Fox, 2014). However, polymorphisms that persist across species (Jamie & Meier, 2020) represent evolutionary enigmas. How can polymorphisms transcend species boundaries? Convergent evolution (Gould, 1989; Conway, 2005) including developmental convergence (i.e. independent changes in gene expression), introgression (Mallet, 2005; Nolte et al., 2009), and the maintenance of an ancestral polymorphism (Igić & Kohn, 2001; Igić et al., 2008) are three plausible evolutionary avenues leading to shared polymorphisms among closely related species.

Color polymorphisms are widespread in nature but only a few cases of trans-specific color polymorphisms have been reported, and mostly restricted to animals (reviewed in McKinnon & Pierotti, 2010; Martin et al., 2013; McLean & Stuart-Fox, 2014; McRobie et al., 2019). Few examples have been investigated in plants. The closest situation for flower color is the repeated transitions from blue to red flowers (Wessinger & Rausher, 2013) associated with shifts from bee- to hummingbird-pollination (Thompson & Wilson, 2008; Muchhala et al., 2014; Wessinger et al., 2019) across a diversity of angiosperm lineages. However, in this case intraspecfic blue-red flower color polymorphisms (variation within populations) are very rare since either (1) multiple mutations are required to accomplish this shift and/or (2) the color is associated with a pollinator shift leading to reproductive isolation between the color types thereby fixing the color differences in newly formed species (but see Stankowski and Streisfeld 2015).

Unlike the blue to red flower color transitions leading to speciation, the blue-orange flower color polymorphism in the closely related species *Lysimachia arvensis* and *L. monelli* (*La* and *Lm* herein, Figure 1A) does not confer a pollinator shift (Jiménez-López et al., 2019; Sánchez-Cabrera, 2023). Instead, in *La* the color polymorphism is driven by abiotic non-pollinator agents of selection such as drought and sunlight intensity (Arista et al., 2013, Ortiz et al., 2015) and temperature (Jiménez-López et al., 2023; B. Carter et al., unpublished data) where blue flowered individuals are fitter than orange ones in drier, sunnier and hotter environments. *La*, is an annual native to the Mediterranean Basin and central and northern Europe. A recent autopolyploid origin has been suggested for *La* for three reasons: (1) due to the lack of ITS polymorphisms (Jiménez-López et al., 2022), (2) the presence of four identical copies of one set of chromosomes (Monein et al., 2003), (3) and the phylogenetic configuration of the group (Jiménez-López et al., 2022). Color morph frequencies range from 0% to 100% with a steep cline in central-southern Mediterranean region where mixed populations are common (e.g., Portugal, Spain, Italy and France; Arista et al., 2013). The cline and resulting association with abiotic forces strongly argue against pollinator mediated selection and instead point to abiotic factors which have been substantiated in a greenhouse study (Arista et al., 2013). In this species, the shift from blue to orange (Sánchez-Cabrera et al., 2021) correlates with a biochemical transition from malvidin 3-rhamnoside to pelargonidin 3-glucoside (Harborne, 1968; Sánchez-Cabrera et al., 2021). Orange flowers have increased expression of *DFR-2*, a duplicate gene only found in orange-flowered individuals with non-synonymous SNPs suggesting substrate specificity for dihydrokaempferol drawing flux down the pelargonidin branch of the anthocyanin biosynthetic pathway (ABP) (Sánchez-Cabrera et al., 2021).

**Figure 1.**
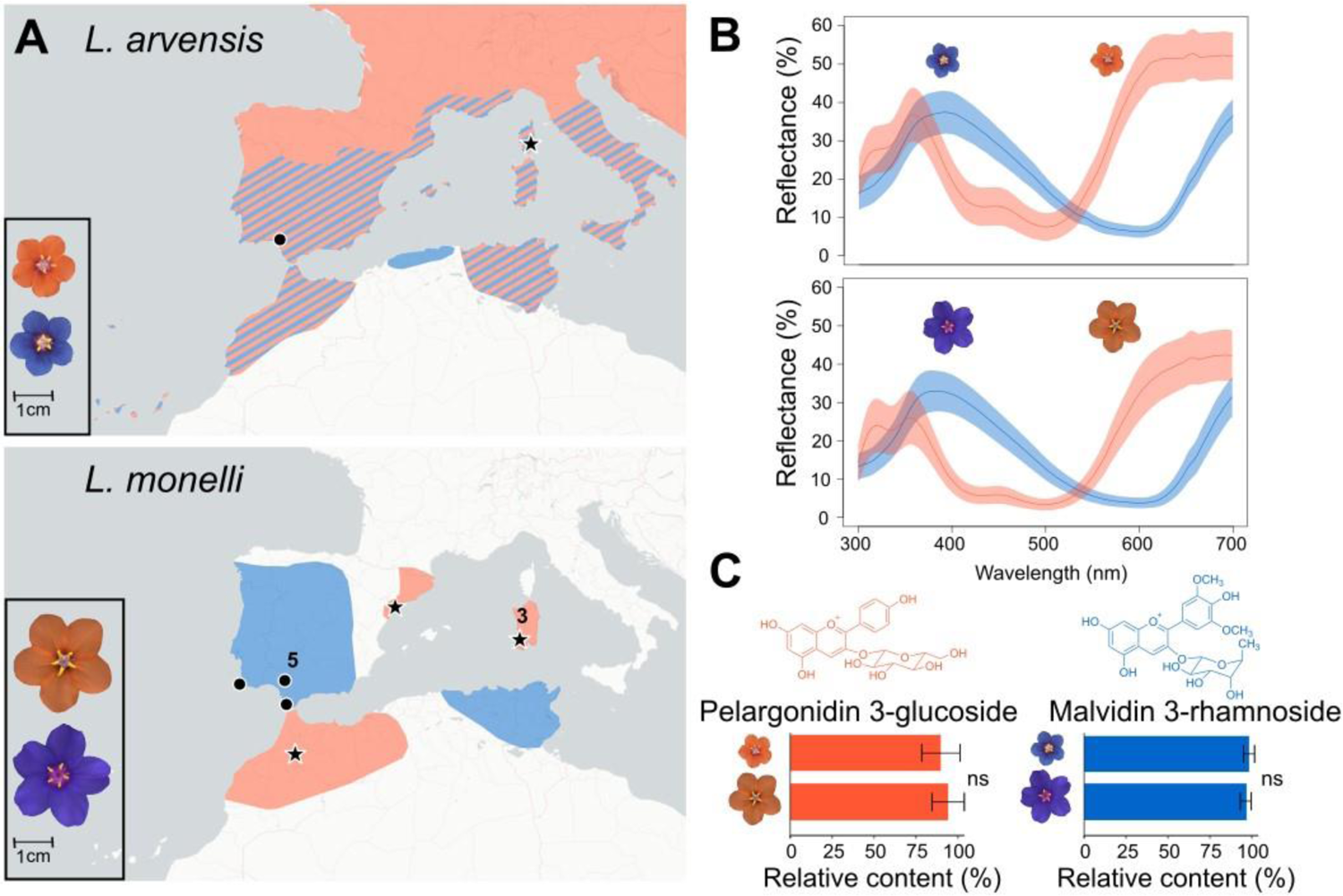
Flower color characterization of blue and orange *Lysimachia arvensis* and *L. monelli*. (A) Scaled images and geographic distributions of the two flower color morphs of both species. Coexistence of the *L. arvensis* color morphs is indicated with diagonal striping. Populations sampled for transcriptome comparisons are indicated with black circles (blue) and black stars (orange) (numbers indicate >1 geographically adjacent populations were sampled for *L. monelli*). (B) UV-vis reflectance spectra of blue and orange petals of both species showing mean curves with 95% confidence intervals in colored shading above and below the mean curve. (C) Molecular structures and relative content from the biochemical analysis of the primary anthocyanins in both species. Non-significant differences (ns) in relative content are indicated using a Mann-Whitney U test. In *L. arvensis*, orange samples include pelargonidin 3-glucoside derivatives. Error bars represent standard error (SEM).

Herein, we investigate the shared blue-orange flower color polymorphism in the closely related *Lm*, a diploid perennial, that co-occurs with *La* in some populations (Figure 1). Blue flowered populations of *Lm* are more common in drier habitats of central and southwestern Iberian Peninsula and eastern North Africa; while orange-flowered populations are more often found in wetter habitats of northeastern Iberian Peninsula, western North Africa, and Sardinia (Gibbs & Talavera, 2001; Sánchez-Cabrera, 2022) (Figure 1A). In *Lm*, no populations have both color morphs (Sánchez-Cabrera, 2023).

To determine the origin and maintenance of the shared blue-orange flower color variation in this species of *Lysimachia* and how it relates to the strikingly similar blue-orange flower color variation in *La*, we investigate the similarities in petal reflectance spectra, flavonoid biochemistry and compare petal transcriptomes allowing us to weigh four possible hypotheses explaining the origin and persistence of this flower color polymorphism: convergence, introgression, ancestral polymorphism or the non-monophyly of species (Figure 2).

**Figure 2.**
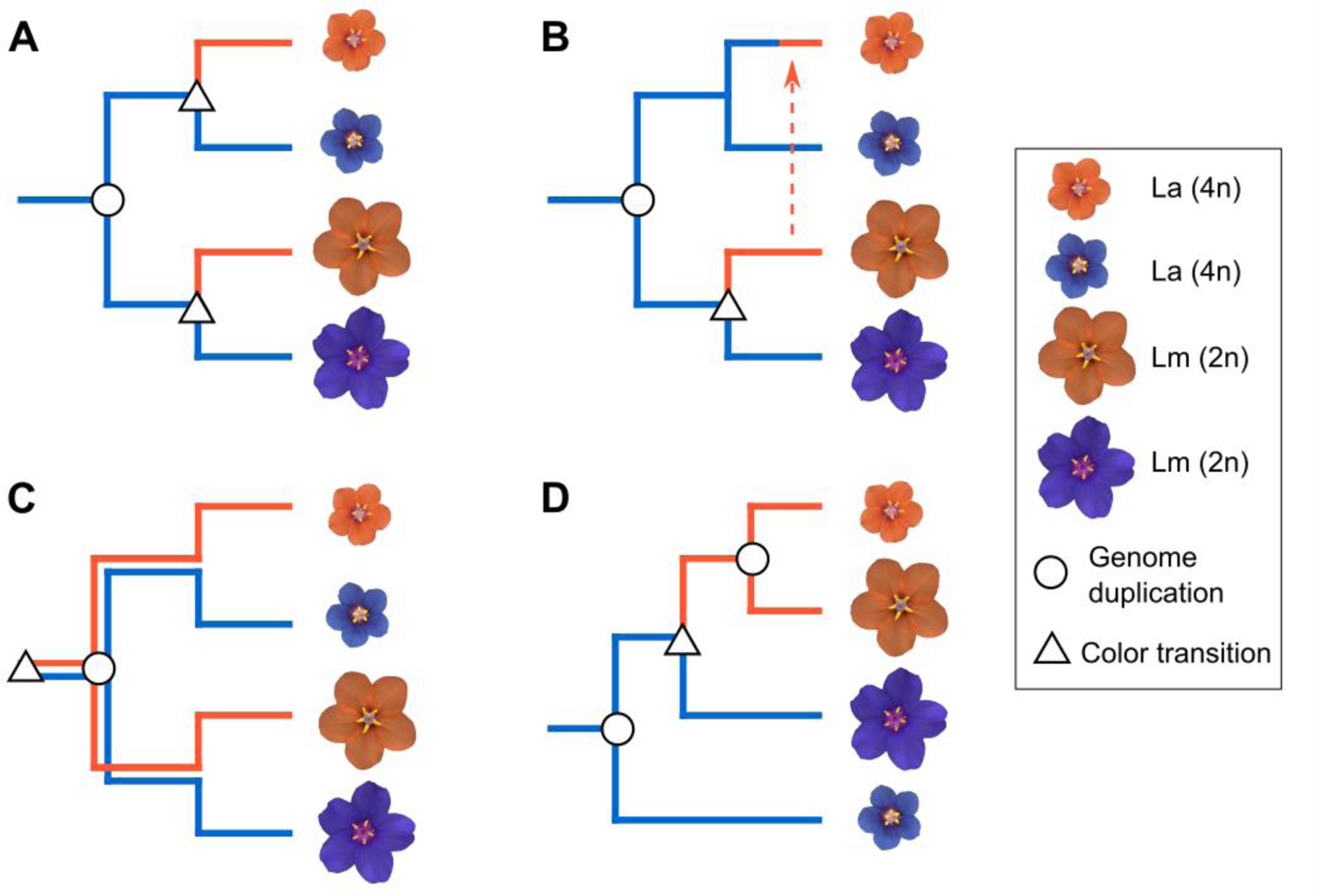
Four evolutionary hypotheses of petal color evolution in *L. arvensis* and *L. monelli*. Assuming the ancestral state is blue and diploid, the most parsimonious color shifts are indicated by a triangle and ploidy changes are indicated with a circle (placed at the node since they likely drive divergence). In (A), convergent evolution of orange petals is coupled with a single origin of the tetraploid lineage predicts independent and distinct molecular causes for the color change in each lineage. In (B), a single origin of orange petals in *L. monelli* followed by introgression to *L. arvensis* requires only one shift to orange and one polyploid event but requires the lineages to be able to hybridize. In (C), a single origin of orange in the common ancestor creates an ancestral polymorphism that transcends the speciation event and persists to the present in both species, requiring one polyploidy event. In (D), non-monophyletic species allow for a single origin of orange petals but requires independent origins of the tetraploid lineages.

## Results

### Petal UV-Vis Spectra

Reflectance spectra of petals for both species are very similar across the UV and visible wavelengths when comparing the same color morphs, yet quite different between color morphs (Figure 1B; Appendix 1—figure 1). Spectra distances between color morphs are ∼7x larger than spectra distances within morphs between species (Permutational Manova, R2 = 0.75 vs. 0.15; Appendix 1—figure 1). Orange petals have double UV reflectance peaks that are absent in blue petals and differ in their primary inflection points (380, 570nm for orange and 480, 660nm for blue).

### Petal Biochemical Comparisons

Biochemical analyses of petal extracts in acidified methanol using HPLC-MS-DAD revealed the underlying biochemical differences between blue and orange morphs are nearly identical between the two species. Blue petals of both species are composed of malvidin 3-rhamnoside, however small amounts of the aglycones malvidin and delphinidin, and a flavonol derivative were also detected in both species (Figure 1C; Appendix 1—table 1; Sánchez-Cabrera et al., 2021). In contrast, orange petals of both species accumulate principally pelargonidin 3-glucoside and to a lesser degree, other pelargonidin derivatives. There were no significant differences in the relative content of malvidin 3-rhamnoside and pelargonidin 3-glucoside derivatives when comparing the blue and orange samples of these two species, respectively (Figure 1C).

**Table 1.**
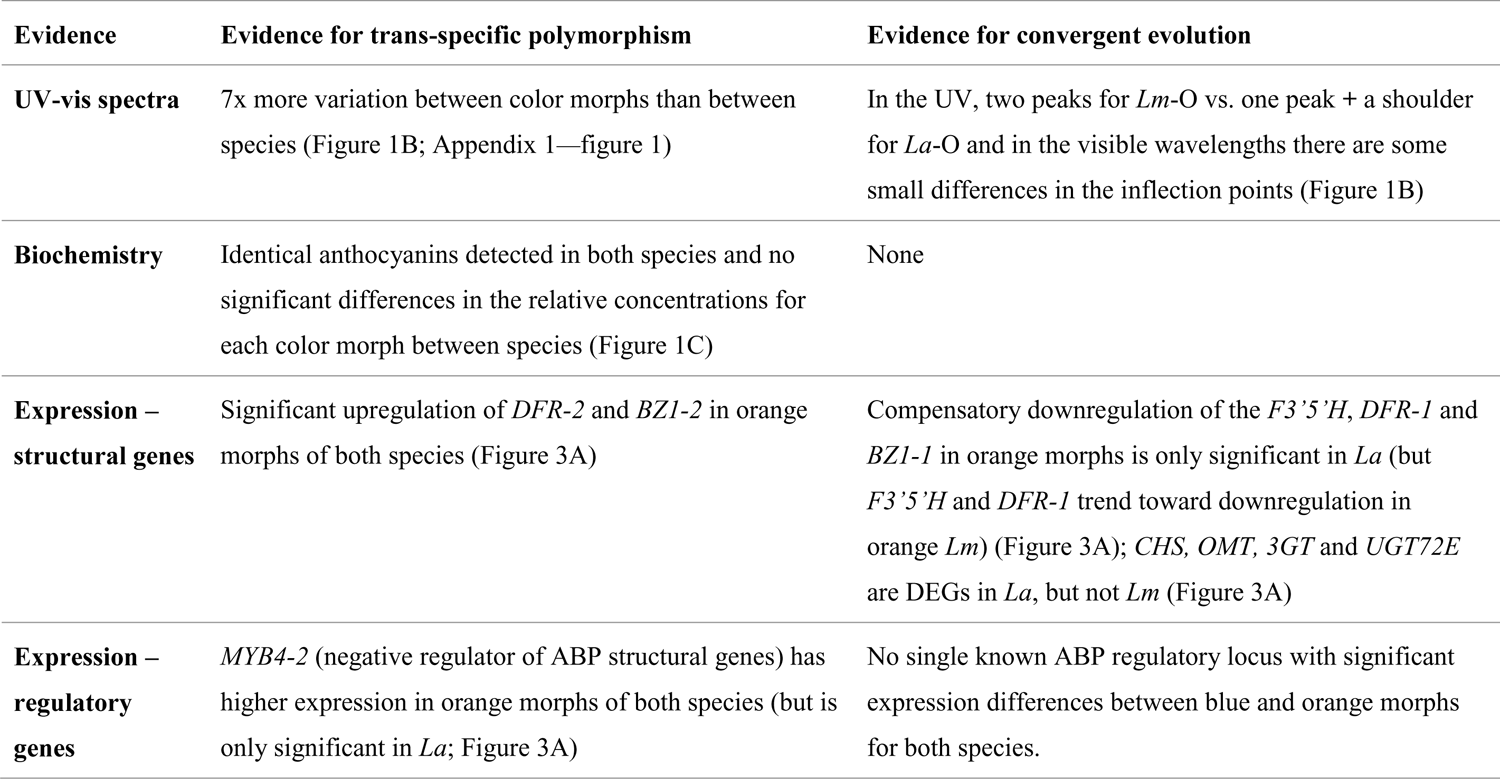
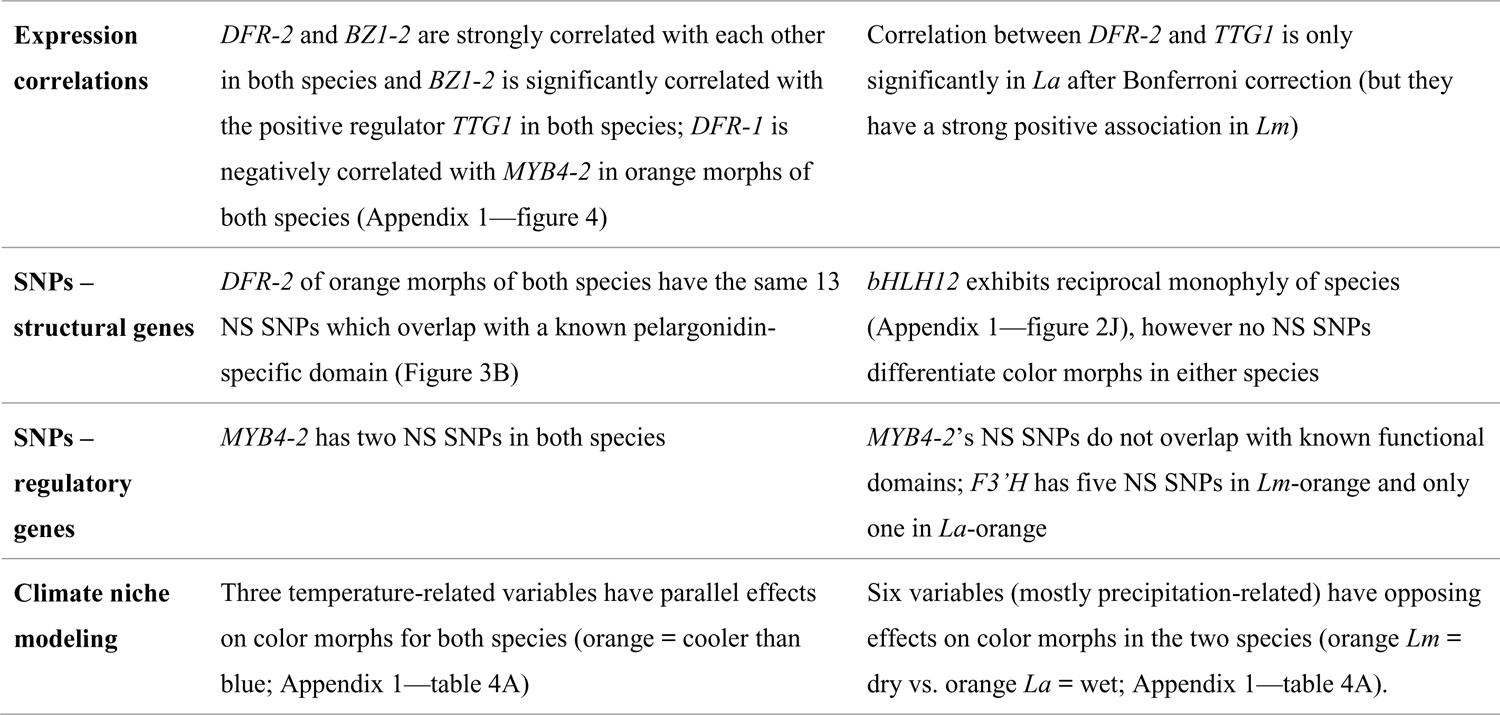
Evidence for a single origin versus convergent evolution of the blue to orange flower color shift in *Lm* and *La*.

### Transcriptome comparisons

#### Differential expression

Transcriptome results for *Lm* petals identified 37,552 distinct genes (see Sánchez-Cabrera et al., 2021 for *La* results including the absence of any evidence of divergent homologous copies of these genes). Of those, 336 were differentially expressed between blue and orange flowers (1.19x more genes with O > B expression; Chi-Square = 14307; p = < 2.2e-16). A total of 128 flavonoid biosynthetic pathway (FBP) genes were detected, although only five had significant differential expression, four of which had O > B expression in *Lm* (Appendix 1—table 2), three of which are in the core ABP (Figure 3C)

**Figure 3.**
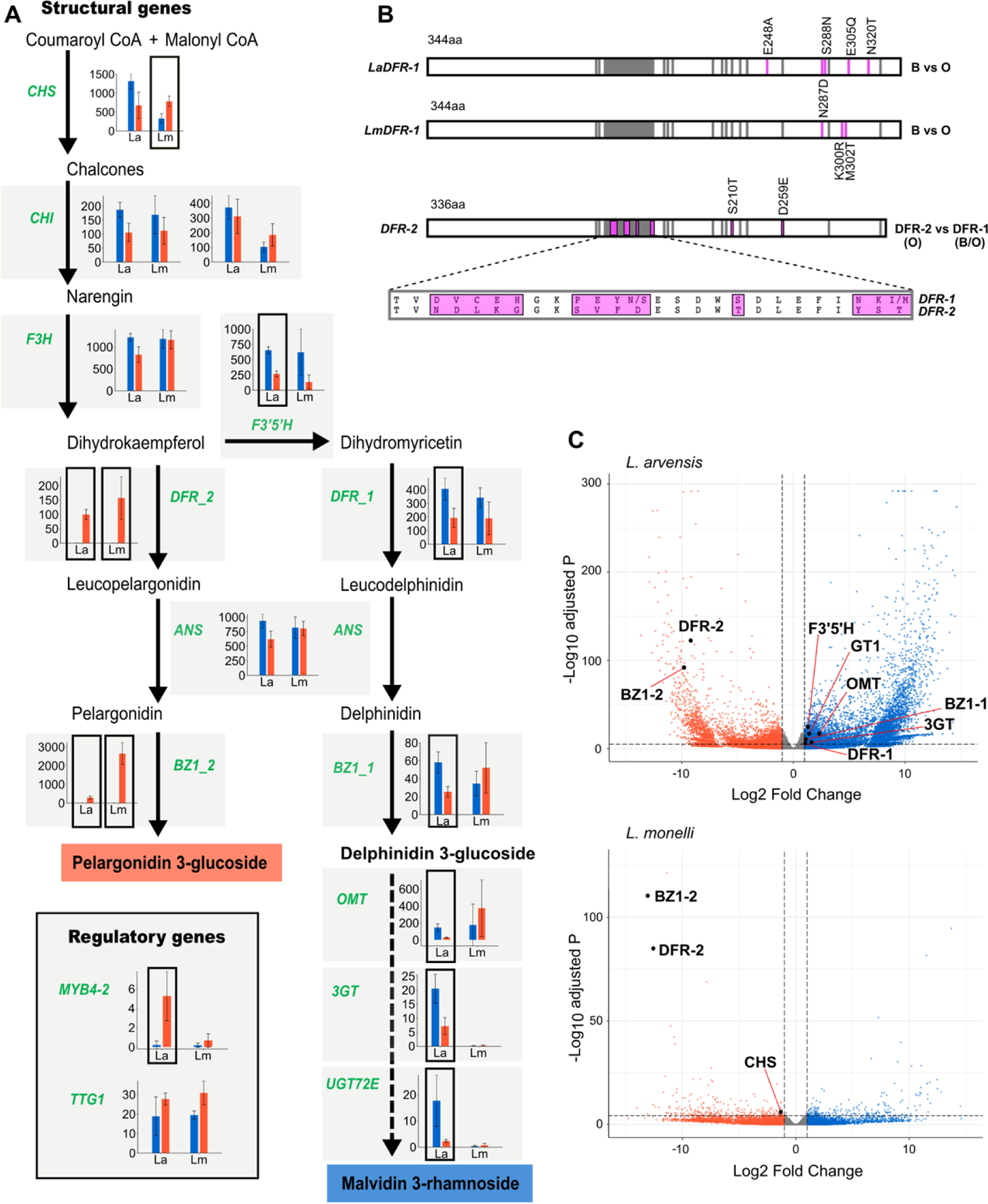
Gene expression comparisons of blue and orange morphs of two species of *Lysimachia*. (A) Suggested Anthocyanin Biosynthetic Pathway (ABP) producing blue (malvidin 3-rhamnoside) and orange (pelargonidin 3-glucoside) flower colors in these two species. Mean expression (TMM +/- standard error) in both species (La and Lm) are shown for each ABP gene (green font). Significantly differentially expressed genes (DEGs) are indicated with a black box around the bar plots. (B) Diagrams of the *DFR* coding sequence showing color-differentiating non-synonymous SNPs in pink (>70% frequency difference) between blue and orange *DFR-1* of *L. arvensis* and *L. monelli*, and differences between *DFR-1* (blue and orange) and *DFR-2* (orange) of both species. Gray areas indicate substrate specificity and active sites of *DFR* genes. The magnified region comparing *DFR-1* and *DFR-2* shows many NS SNPs (pink) in the largest substrate specificity region. (C) Volcano plot comparing the entire transcriptomes of *L. arvensis* and *L. monelli* highlighting ABP genes with significant differential expression.

In comparing FBP gene expression in *Lm* to *La,* three genes (*DFR-2*, *BZ1-2* and *Caffeoyl CoA-1*) were significantly differentially expressed in both species (FDR<10^-5^ and log_2_FC>1; Figure 3C; Appendix 1—table 2; see Sánchez-Cabrera et al., 2021). Two key ABP genes had high expression in orange petals and were nearly undetectable in blue petals (Figure 3A, C). First, *DFR-2* with substrate specificity for pelargonidin (see references in Sánchez-Cabrera et al., 2021) had high expression in orange petals (*LmDFR-2* orange mean = 157.18; *LaDFR-2* orange mean = 99.80; Figure 3A). The second is *BZ1-2*, a gene responsible for glycosylation that stabilizes the orange pigment (*LmBZ1-2* orange mean = 2651.84; *LaBZ1-2* orange mean = 294.55; Figure 3A).

However, the reciprocal decrease in expression for the primarily blue paralogue (*DFR-1* and *BZ1-1*) in orange flowers was only significant in *La*, but trends in the same direction at similar magnitude for *LmDFR-1* (blue is ∼2x higher than orange) (Figure 3A). The third gene, *Caffeoyl CoA-1*, is an early gene in the ABP that functions before the first dedicated step in the ABP (not previously reported to be involved in this type of color change; Appendix 1—figure 2B). It had higher expression in blue petals of both species and was nearly undetectable in orange petals of both species - the opposite pattern as that described above for *DFR-2* and *BZ1-2*. In addition to not having known function in the biochemistry of blue and orange, this gene is also unlikely responsible for the color change because this copy of Caffeoyl CoA has the lowest expression of the three copies expressed in petals (Appendix 1—figure 2B).

Some additional ABP genes likely involved in this color shift showed significant differential expression between colors of *La* and consistent, yet non-significant trends, in *Lm*. For example, *DFR-1* and *F3’5’H* had higher expression in blue petals than in orange petals in both species, but was only significant in *La* (2.10x and 2.45x more in *La*B and 1.80x and 4.71x more in *Lm*B, respectively; Figure 3A). Another example of an ABP gene with significant DEG in *La* and a similar trend, only significant with a Mann-Whitney U test, in *Lm* is *MYB4-2* which has higher expression in orange petals compared to blue petals (25.75x more in *La*O and 3.83x more in *Lm*O; Figure 3A).

Alternatively, *CHS* showed significant differential expression between colors of *Lm*, but not significant nor consistent in *La* (1.95x more in *La*B, but 2.36x more in *Lm*O; Figure 3A) and is unlikely involved in the transition from blue to orange. Finally, a third *DFR* (*DFR-3*; Figure 4) was detected in both species, but exhibited very low expression (maximum TMM = 1.07) indicating it is not the primary gene copy at this step in the ABP in petals (*DFR-1* and *DFR-2* have >100x higher expression).

**Figure 4.**
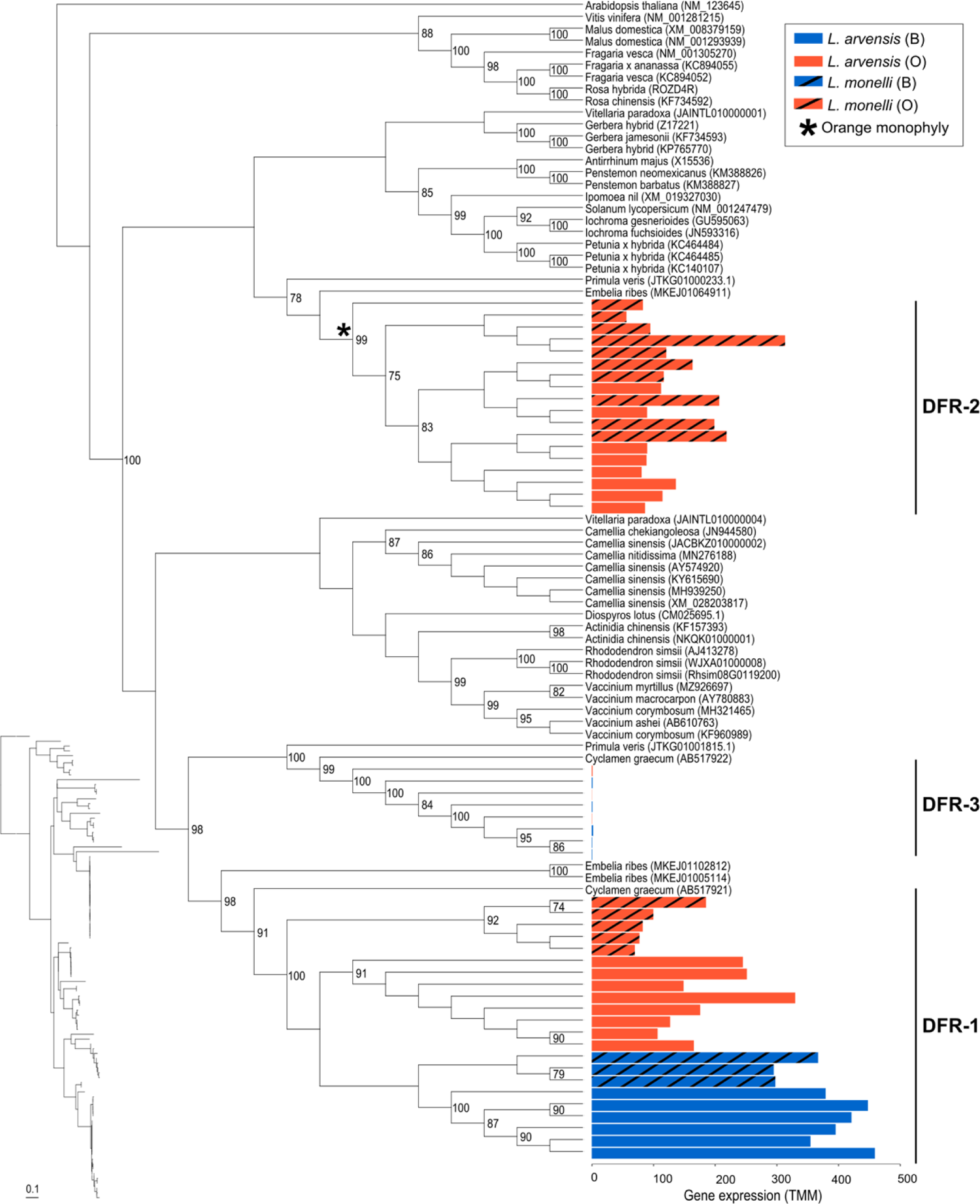
Maximum likelihood cladogram of the coding sequences of three *DFR* paralogues expressed in two *Lysimachia* species. Outgroups were selected from top BLASTn hits and, when available, from the genomes of closely related species (*Camelia sinensis*, *Embelia ribes* and *Primula veris*). Outgroups are indicated by scientific name followed by Genbank Accession numbers. Bootstrap values are provided to the right of the nodes when greater than 70%. The bar plot shows the expression level (TMM) for blue and orange flowers of *L. arvensis* (solid) and *L. monelli* (black stripes). The monophyletic clade of orange petal samples for *DFR-2* is indicated with an asterisk (*). A phylogram (inset) is provided with the samples in the same order as the cladogram to compare relative branch lengths. Scale bar is in substitutions per site.

#### Phylogenetic analysis of ABP sequences

We compared coding sequences of a total of 22 ABP genes that were either differentially expressed in one of the species or potential candidate genes to determine whether species were monophyletic (convergence) or if color was monophyletic and transcended speciation (ancestral polymorphism). Individual phylogenetic analyses for all 22 ABP genes (10 structural and 12 regulatory) consistently recovered a monophyletic clade of *Lysimachia* samples suggesting these were broadly orthologous comparisons (>70% bootstrap support; Appendix 1—table 3 and figure 2). For two key ABP genes in the transition from blue to orange (*DFR-2* and *BZ1-2*), all orange samples of both species are strongly supported as monophyletic (99% and 100% bootstrap, respectively; Figure 4; Appendix 1—table 3 and figure 2A). Furthermore, the regulatory locus, *bHLH12* is reciprocally monophyletic for species and within species color morphs (Appendix 1—figure 2J). In a combined phylogenetic analysis of all ABP loci, we recovered monophyletic species and monophyletic colors within each species except for paraphyletic *Lm*O (Appendix 1—figure 3).

#### SNP analysis

To determine if there is an association between genotype and phenotype, we examined SNPs across the FBP loci for all samples. Genes responsible for color differences in one or both species could have non-synonymous (NS) SNPs in functional regions that correlate with color within species if causal changes are in coding regions. Further, if these are shared between species this would provide further evidence supporting a single origin (ancestral polymorphism). We detected the exact same 13 NS SNPs in *DFR-2* of orange *Lm* that differentiate it from *DFR-1* which overlap with known functional regions providing substrate specificity for pelargonidin (Figure 3B) as we detected previously in *La* (see references in Sánchez-Cabrera et al., 2021).

Broadening our analysis to all FBP/ABP genes examined, only two loci have completely fixed differences between color morphs in both species (one NS SNP in *DFR-1* and two NS SNPs in *MYB4-2*; Appendix 1—table 4), but none are located in known functional sites.

By comparing NS SNPs that differentiate the color morphs with the NS SNPs that differentiate the two species, we can begin to hone in on the loci responsible for the color change to eventually address if it was a single event (ancestral polymorphism or introgression) or more than one event (convergence). We find 16 NS SNPs that differentiate the two species in eight genes that we don’t predict are causing the color change (Appendix 1—table 4). Noticeably absent from this list are the loci we have evidence are involved in the color shift (*DFR-1*, *DFR-2*, *BZ1-2*, *MYB 4-2*), and only one NS SNP in the extreme 5’ region of *F3’5’H* (Appendix 1—table 4). Only *F3’H* showed color-specific NS SNPs, higher in *Lm* than in *La* (five vs. one), however there are even more NS SNPs between the species (seven) (Appendix 1—table 4).

### Structural and regulatory gene interaction

To help discern if the shared expression differences in orange petals of the two species for *DFR-2* and *BZ1-2* are controlled by the same or different regulatory genes, we looked for patterns of correlated expression with the MYB-bHLH-WD40 regulatory complex. The rationale for investigating expression correlations is that if color has evolved via independent changes in expression (i.e. developmental convergence), then we expect gene expression correlations in the gene involved to be species specific.

Using a hypothesis testing approach (alpha = 0.05) in both species, *DFR-2* and *BZ1-2* were significantly positively correlated with one another (R = 0.73, p = 8.8e-05 in *La*; R = 0.62 and p = 2.9e-04 in *Lm*) and with *TTG1*, a WD40 known to activate ABP structural genes (across both genes and species p < 0.0051; Appendix 1—figure 4). Furthermore, *DFR-1* expression is negatively correlated with *MYB4-2* expression, a known repressor of ABP structural genes (R = −0.68 and p = 2.5e-04 in *La*; R = −0.62 and p = 1.4e-04 in *Lm*; Appendix 1—figure 4). *MYB4-2* is positively correlated with *DFR-2* (R = 0.39 and p = 0.37 in *La*; R = 0.47 and p = 0.0062 in *Lm*), but non-significant after Bonferroni correction.

### Reproductive isolation between *La* and *Lm*

None of the crosses performed between *La* and *Lm* produced fruits irrespective of crossing direction and flower color (n=53 pollinations, see Appendix 1—table 5). In contrast, morphs within species were interfertile. Between morph crosses in *Lm* produced a mean fruit-set of 47.86% ± 0.39% (n=575) and a mean number of seeds per fruit of 16.25 ± 9.16 (n=287) (Sánchez-Cabrera, 2023). Whereas in *La*, 100% of between morph crosses produced fruits containing 16.95 ± 7.19 seeds per fruit (n= 97) (Jiménez-López et al., 2023).

### Climate niche modeling

The four morphs were statistically different from one another across most of the 19 BIOCLIM variables examined (Appendix 1—table 6). The orange and blue *La* morphs differed significantly from each other for 17 of the 19 climate variables, and the two *Lm* morphs differed for nine variables. The two orange morphs differed from one another for 16 variables and the two blue morphs differed across 11 variables.

Nine variables were significantly different between the two blue morphs and between the two orange morphs of both species. Of these, two temperature variables differed in parallel (e.g., for “isothermality” orange morphs showed lower values than blue morphs for both species) (Appendix 1—figure 5A), while three temperature and four precipitation variables differed in opposing directions (e.g., for “annual precipitation” orange *La* were wetter than blue *La*, but orange *Lm* were drier than blue *Lm*) (Appendix 1—figure 5C).

The logistic regression results showed that the two best models using AIC did not employ any of the same variables. The best model for *La* included (in decreasing order of importance based on standardized coefficients) precipitation seasonality, isothermality and temperature seasonality, while the model for *Lm* consisted of precipitation of the wettest month and mean temperature of the wettest quarter (Appendix 1—table 7).

Since the ancestral state in *Lysimachia* is blue flowers, we hypothesized that blue would be less likely to diverge in their climate niche between the two species compared to orange morphs of the two species. We tested this with a Monte Carlo procedure that randomized colors within species and then compared observed climatic differences with differences from the null distributions. Across the 19 BIOCLIM variables, blues were more divergent between the two species than expected by chance for only one variable while orange samples were more divergent between the two species for 15 variables (∼79% of variables studied), and neither color morph was significantly different for three variables (Appendix 1—table 8).

## Discussion

There are many similarities (and some differences) between color morphs of *L. arvensis* and *L. monelli* that can be used to assess the alternative hypotheses (Table 1), however definitive evidence excluding all but one of these hypotheses remains elusive. If orange evolved independently in each species (i.e. convergence; Figure 2A), we would expect distinct molecular causes (Streisfeld & Rausher, 2009; Larter et al. 2018; Ng & Smith, 2016), which we don’t have any strong evidence for in these two species of *Lysimachia*, but the exact mutation(s) responsible for the color shift in each species awaits further genetic dissection. If the remarkably similar upregulation of *DFR-2* was activated by different MYBs in orange petals of these two species, we would expect distinct correlations in expression. Instead, *DFR-2* and *MYB4-2* are positively correlated in both species (but not significant after Bonferroni correction; Figure S5). Alternatively, the likelihood of a single shift from blue to orange followed by introgression into the other species (Figure 2B) as clearly documented in the *Diplacus* (*Mimulus*) *aurantiacus* complex (Stankowski and Streisfeld 2015, Short and Streisfeld 2023) is unlikely in *Lysimachia* for two reasons. First, although the two species co-occur, hybrids with intermediate flower size and plant size have never been observed (Pujadas, 1997) unlike in the *Diplacus aurantiacus* complex where hybrid zones are well documented. Second, numerous attempted crosses of the two species reveal complete reproductive isolation (Table S7), presumable because of the ploidy differences – whereas the members of the *Diplacus aurantiacus* complex are at least partially interfertile (Stankowski and Streisfeld 2015). But, we can only test for the likelihood of genetic exchanges between the two species in the present – there is always the possibility that there was some gene exchange soon after the (auto)polyploid speciation event. Even in that scenario, the single origin followed by introgression would require the long-term persistence of this polymorphism even though it may not have transcended the speciation event.

Given the evidence presented herein, the most parsimonious explanation starts with a single origin of orange from blue in a common ancestor of *La* and *Lm* - creating an ancestral polymorphism that then transcends the speciation event separating these two closely related species (Figure 2C). In this case, both blue and orange individuals would have persisted through the speciation process including polyploidy, loss of self-incompatibility, significant reduction in flower size, and a shift from perennial to annual. Although the persistence of polymorphisms across species has been documented in a few exceptional animals (see Jamie & Meier, 2020 for a review), this is the first example in plants with regards to a flower color polymorphism, however an analogous pattern of trans-specific, shared polymorphism is well documented for self-incompatibility alleles (Igić et al. 2008). Furthermore, this flower color example appears to be driven by non-pollinator agents of selection – specifically the distinct climatic niches in each species, especially those involving precipitation variables (Arista et al., 2013; Ortiz et al., 2015; Sánchez-Cabrera, 2023), yet does not appear to be maintained by negative frequency dependent selection, the most common mechanism maintaining polymorphisms over long periods of time in other species of plants and animals.

A final alternative to ancestral polymorphism could involve a more complex evolutionary history where these two species are not reciprocally monophyletic (Jiménez-López et al., 2022; Figure 2D). This is unlikely given the combined phylogenetic analysis of 22 ABP loci which strongly supports the monophyly of both species (Appendix 1—figure 3). Moreover, in this scenario, a single origin of orange would have to be followed by repeated origins of polyploidy, independent losses of self-incompatibility, recurrent reduction in flower size and separate shifts to the annual habit which is substantially less parsimonious than the persistence of an ancestral polymorphism (Figure 2C vs. D).

The color morphs of these two *Lysimachia* species have nearly identical biochemical and molecular underpinnings as measured thus far. The evolution of orange pelargonidin 3-glucoside from blue malvidin 3-rhamnoside requires (1) redirecting ABP flux down the pelargonidin branch and (2) stabilizing the newly formed orange anthocyanidin with glucose instead of rhamnose. Both steps are accomplished in the same way in both species - via the recruitment of orange-specific paralogues that are undetectable in transcriptomes of blue petals. In fact, the copy of *DFR* unique to orange petals (*DFR-2*) has nearly the same overall gene expression levels as the shared copy (*DFR-1*), but its expression is undetected in blue petals of both species. Furthermore, *DFR-2* of both species contain the same 13 NS SNPs in the substrate specificity region that differentiate it from *DFR-1*, which likely allows it to outcompete *F3’5’H* (and *F3’H*) for dihydrokaempferol thereby shunting flux down the pelargonidin branch leading to orange petals (Johnson et al., 2001; Petit et al., 2007; Katsu et al., 2017). The substantial genetic distance between *DFR-1* and *DFR-2* suggests that this duplication predated the genus and maybe even the Primulaceae family – a molecular toolkit deployed when pelargonidin provides a selective advantage over malvidin. Although the exact regulatory locus (loci) responsible for these shifts in expression will require further genetic analysis, we hypothesize it involves *TTG1* (pelargonidin-specific activator) and *MYB4-2* (malvidin-specific negative regulator and potential positive activator of *DFR-2*). We find no evidence of independent NS SNPs in either *TTG1*, nor *MYB4-2*, suggesting that the regulatory control of this flower color polymorphism has only evolved once. Instead, we see remarkably similar expression patterns and even some shared non-synonymous SNPs in both regulatory loci that are the most likely candidates for the molecular basis for this color shift.

The coordinated changes in ABP gene expression is like other blue to red flower color shifts that involve the recruitment of substrate specific *DFR* copies with correlated downregulation of the alternative side-branch gene expression (e.g., *F3’H* and *F3’5’H*) (Zufall & Rausher, 2004; Des Marais & Rausher, 2010). We also find decreased expression in *F3’5’H* (but not *F3’H*) in orange petals of both species, however, this is only significant in *La* due to high expression variation among the blue *Lm* samples (Figure 3). Regardless, the shared expression and nearly identical coding sequences of orange *DFR-2* in *La* and *Lm* are strong evidence of a single origin that predates this speciation event and has persisted to the present in these distinct lineages.

Similarly, orange petals employ an orange-specific glycosyltransferase (*BZ1-2*), undetected in blue petals, glycosylating the orange pigment in the same way in both species. The shift from blue to orange is correlated with a dramatic increase in expression of the glycosyltransferase *BZ1-2* (similar to the *DFR*s mentioned above), which is likely involved in the stability, solubility, storage and biological activity of this particular anthocyanin (Yonekura-Sakakibara et al., 2008). There are no NS SNPs between orange samples of *La* and *Lm* (same evolutionary history), however two *BZ1* paralogues are highly divergent (>60% nucleotide divergence) and unalignable indicating a relatively ancient duplication event). The orange-specific *BZ1-2* of both species contains NS SNPs correlating with glucose-specificity as characterized in *Vitis vinifera* and *Medicago truncata* (He et al., 2006; Wang, 2009). In particular, there are 20 NS SNPs distinguishing these two paralogues in the PSPG-box, a conserved 44 amino acid region found in all plant UFGTs (Caputi et al., 2012) that likely confers glucose-specificity of *BZ1-2*. We infer that the two paralogues likely have different functions regarding which sugar they add and their efficiency in doing so (Sun et al., 2016), but within a color morph for both species, their sequence similarity suggest they perform the same function. In contrast to *BZ1-2*, *BZ1-1* has variable expression in both color morphs - *La* has significantly higher expression in blue than orange as expected, but mean expression is similar in blue and orange *Lm* (and not significant). Regardless of whether there is compensatory downregulation in orange petals of *BZ1-1*, we know that *BZ1-2* has 10-50x higher expression in orange and we predict that it has higher efficiency in adding glucose to pelargonidin of orange petals than *BZ1-1* (Sun et al., 2016). Although there are other examples of closely related species exhibiting the same flower color because they accumulated similar major categories of anthocyanidins (i.e. aglycones) (e.g. Ng & Smith, 2016; Larter et al., 2018; Ogutcen et al., 2020), the specific anthocyanins they accumulate are generally distinct due to the enormous diversity of biochemical decorations in the flavonoids (Passeri et al., 2016; Berardi et al., 2021). In these two *Lysimachia* species, orange morphs use the same *3GT* correlated with the predominant glycosylation of pelargonidin clearly indicating a single ancestral evolutionary transition shared by both species that has transcended the speciation event.

If *DFR-2* and *BZ1-2* are similarly upregulated in orange *La* and *Lm*, we predict they will be controlled by the same regulatory gene(s) in both species if there was a single ancestral polymorphism. In fact, expression of *DFR-2* and *BZ1-2* in orange petals of both species is positively correlated with the same regulatory gene *TTG1* (WD40) known to control these two ABP structural genes (Tiang & Wang, 2020) (Appendix 1— figure 4). Looking at the sequence of this regulatory gene, there is only one color differentiating NS SNP in *La* and none in *Lm* strongly suggesting another trans-acting regulatory gene is likely responsible for the differential expression of *TTG1* in blue and orange morphs of these two *Lysimachia* species. However, *MYB4-2* has two NS SNPs shared by both species that positively correlates with *DFR-2* expression (Appendix 1— figure 4).

For both *Lysimachia* species, *DFR-1* expression in orange petals is negatively correlated with *MYB4-2* expression (Appendix 1—figure 4). Moreover, in *La* this increase of *MYB4-2* is correlated with a decrease in *F3’5’H* in orange petals, facilitating the redirection of ABP flux to the pelargonidin branch of the pathway. Although *MYB4-2* has low expression in both morphs of *Lm* and is not significantly correlated with *DFR-1* nor *F3’5’H*, it exhibited significant differential expression between morphs (Mann-Whitney U test, U = 19, p = 0.021). In *Arabidopsis thaliana* and *Vitis vinifera*, *MYB4* represses anthocyanin biosynthesis either through direct binding to the promoter regions of ABP genes (*ANS*, *DFR* and *UFGT*) or by displacing the activator MYB in the MBW complex (Pérez-Díaz et al., 2016; Chen et al., 2019). However, how could *MYB4-2* repress *DFR-1* and not *DFR-2*? Like in *Lotus* (Yoshida et al., 2010) and *Fagopyrum* (Katsu et al., 2017) where different *DFR* paralogues are independently regulated by their unique promoter sequences, we predict that *MYB4-2* has differential regulatory effects on the two primary *DFR* paralogues. Overall, we argue that *MYB4-2* contributes to redirecting ABP flux down the pelargonidin branch of the pathway by negatively regulating the non-orange-specific paralogues at the critical branchpoint (*DFR-1* and *F3’5’H*) in both species similarly.

Previous experimental work and our climate niche modelling indicate that there may be pleiotropic effects of petal color. Orange morphs of both species are found in locations with colder winters than blue morphs (Appendix 1—figure 5A). However, the two morphs appear to respond in opposing directions to primarily moisture-related climate niche variables (Appendix 1—figure 5C). Orange *Lm* is found in habitats with a wide range of precipitation while orange *La* is found in wet habitats. Although the climate niche of the blue morphs of the two species are often distinct, the orange samples are clearly driving these orthogonal responses to precipitation. Why a biochemically and genetically similar polymorphism shared between different species would have contrasting ecological side-effects remains unclear. Previous experimental work in *La* shows that blue-flowered individuals (containing rhamnose stabilized malvidin) inhabit environments with lower precipitation and higher solar radiation than orange-flowered plants with glucose-bound pelargonidin, potentially linking abiotic stress with differential glycosylation (Arista et al., 2013; Sánchez-Cabrera, 2022). If *BZ1-2* is only found in orange petals and is glucose-specific, then does the type of sugar confer physiological or ecologically-relevant adaptations in *Lysimachia*? The coupling of rhamnose to malvidin found in *Lysimachia* also explains the blue color of *Petunia hybrida* (Kroon et al., 1994), *Lobelia erinus* (Hsu et al., 2017) and *Parochetus communis* (Tatsuzawa, 2020) and is known to provide stress tolerance (e.g. UV response) when rhamnose is bound to flavonoids (Li et al., 2017; Jiang et al., 2021).

However, in *Lm*, the opposite is true for several climate niche parameters, especially with regard to precipitation (Appendix 1—table 7 and figure 5C). It appears this shared color polymorphism that may have transcended speciation confers unique climate niches.

Further evidence of gene function, enzyme kinetics and genetic association must be conducted before any final conclusions can be made regarding the molecular causes and ecological consequences of this flower color variation. However, the data presented herein lean toward a single biochemical and molecular footprint shared by both species. If the shift from blue to orange happened twice (i.e. convergence), it would be remarkable since it would most likely be in the same trans-acting regulator of *TTG1*.

Additional experimental approaches (genetic dissection, transformation, knock-out, *DFR* specificity enzyme assays, etc.) promise to identify the type, number and order of mutations responsible for these flower color changes, yet given the phenotypic, biochemical and transcriptome similarities described here, we have yet to find any strong evidence rejecting the ancestral polymorphism hypothesis.

## Methods

### Population sampling

We collected stem cuttings from blue-flowered populations from southern Spain (seven of *Lm* and one of *La*) and from orange-flowered populations from northern Spain, Morocco and Italy (five of *Lm* and one of *La*) (Appendix 1—table 9). Cuttings were rooted in an aeroponic cloner under glasshouse conditions. We also sowed seeds from blue-flowered plants from Portugal in glasshouses at the University of Seville. Voucher specimens were deposited in the University of Seville Herbarium (SEV, Appendix 1— table 9).

We sampled all five petals per flower from a mean of 19 flowers per plant (10 - 24 flowers) from ten blue- and ten orange-flowered plants of *Lm*, and a mean of 30 flowers per plant from eight flowered plants of each color in *La* (see Sánchez-Cabrera et al. (2021) for more details on *La* sampling). In all cases the bullseye at the petal base was avoided. All samples were taken from first-day anthesis flowers and immediately, petals were flash frozen and stored at −80°C. For flavonoid profiling, we collected 10 samples per flower color for *Lm* (flowers from different plants of the same population per sample), including those present in the RNA-Seq analysis (Appendix 1—table 9). *La* samples were obtained from 14 blue- and 11 orange-flowered individuals (see Sánchez-Cabrera et al. 2021). For both species, each sample consisted of two to five flowers without the bullseye.

### UV-Vis petal spectra

Reflectance spectra of the adaxial surface of 44 blue and 67 orange petals of *La* and 75 blue and 50 orange petals of *Lm* were captured using a JAZ A1465 double-beam spectrophotometer (Ocean Optics, Florida, USA). The deuterium-tungsten light source provided reflectance between 300 and 700 nm.

We tested the UV-vis spectra for the relative amounts of variation in two tests (1) between color morphs within a species versus (2) between species within a color morph using permutational multivariate analysis (PERMANOVA test) (Maia & White, 2018). For each test, we computed the Euclidean distances based on the difference in reflectance at each wavelength between all pairs of individual spectra and conducted permutational MANOVA tests (Anderson, 2001) to assess statistical differences (9999 permutations) making quantitative comparative statements between the R^2^ values of the two tests (adonis2 in vegan in R). We visualized the distances between the spectra using metaMDS in vegan in R with 500 tries.

### Identification of flavonoid compounds

Flavonoids were extracted by placing petal samples (see Population sampling section; Appendix 1—table 9) in 1.5 mL microfuge tubes containing 500µl of MeOH with 1% HCl and stored at −80°C. Homogenization was performed with a BeadBeater using 3 x 3.2 mm steel balls for 30 seconds, then centrifuged for 10 min to pellet cellular debris. Samples of the supernatant were loaded on a Dionex UltiMate3000 ultra-high pressure liquid chromatography system equipped with a diode array detector and connected to a Thermo Fleet LCQ mass spectrometer (Thermo-Fisher, Waltham, MA USA). Data were analyzed using Xcalibur software (Thermo Fisher Scientific). Flavonoid identification was based on the compound’s retention time, UV-Vis spectra and whenever possible, chromatographic comparisons with authentic standards: quercetin, luteolin, kaempferol, isorhamnetin, malvidin, pelargonidin, cyanidin and delphinidin (Extrasynthese, Genay, France), previously reported for other *Lysimachia* species (see Sánchez-Cabrera et al., 2021). Relative contents (%) of the primary anthocyanins of each color morph sample, based on the total area under the curve, were calculated and used in statistical comparisons between the two species applying a Mann-Whitney U test in R.

### RNA extraction and reads filtering

Collected petals (see Population sampling section) were homogenized using a mortar and pestle. Total RNA was then extracted following the Qiagen RNeasy Plant Mini Kit protocol (Qiagen, Germany) with the addition of PEG 20,000 mol. wt. (550µl, 2%; Gehrig et al. 2000) before the first filtering step. The addition of PEG was essential to achieve reasonable RNA concentrations for library preparation and sequencing. RNA samples were stored at −80°C until further analysis. RNA concentration and purity was initially assessed with a Nanodrop Nd-1000 (ThermoFisher) and agarose gel, and then confirmed with a Bioanalyzer (Agilent, Santa Clara, CA, USA) before sequencing.

Individual libraries were barcoded, multiplexed and sequenced as 150 bp paired-end reads using two lanes on an Illumina Hi-Seq 2000 (Illumina, San Diego, CA, USA) through Novogene (Beijing, China). Raw paired-end Illumina reads were assessed for quality using FastQC (Andrews, 2010) and were processed using Rcorrector v1.0.4 (Song and Florea, 2015) to correct random sequencing errors. Then, reads were trimmed with Trimmomatic v0.39 (Bolger et al., 2014) to remove any read containing bases with Phred scores lower than 20, low quality reads less than 50 bp long, and any adapter or other Illumina-specific sequences that were still present (Appendix 1—table 10).

### Identification of differentially expressed genes (DEGs)

To determine DEGs for *Lm*, we used the *La* transcriptome assembly with known ABP genes identified (Sánchez-Cabrera et al., 2021) as reference for mapping *Lm* RNA-Seq samples with Bowtie2 software (Langmead & Salzberg, 2012). We calculated Trimmed mean of M-values (TMM, mean of log-expression ratios; Robinson & Oshlack, 2010) for each gene using RSEM software (Li & Dewey, 2011). Then, with EdgeR package (Robinson et al., 2009) in R v4.0.0 (R Core Team, 2020) we determined statistically significant differentially expressed genes (DEGs) between blue and orange petal samples, applying the conservative thresholds for DEG identification: the false discovery rate (FDR) less than 10^-5^ and the expression difference threshold was greater than one log_2_ fold-change (log_2_FC), following Sánchez-Cabrera et al. (2021). To keep biologically interesting genes for differential expression analysis, we considered those genes with more than one count per million (CPM) in a minimum of four samples.

Some regulatory genes have very low expressions and magnitudes were unreliable. For those loci, we performed the non-parametric Mann-Whitney U test to test for differential expression.

### Genes and isotigs selection for analysis and correlated expression analysis

We selected 22 ABP color differentiating structural and regulatory genes with differential expression between colors in either both species, just in one species or not differentially expressed but was a copy of a DEG (see Results section). We established a criterion to select one isotig per gene: (1) samples with less than 20% ambiguous nucleotides in the CDS, (2) samples with more than 100 reads mapped to the sequence, (3) samples with the longest CDS, (4) samples with higher expression. Finally, we selected the isotig which was present in a higher number of samples after applying the criteria. We tested for correlated expression among the 22 structural and regulatory genes in both species using Kendall correlations with Bonferroni correction for multiple testing in R.

### Phylogenetic analyses and non-synonymous SNP identification

We conducted phylogenetic analyses on the selected isotigs for all ABP structural and regulatory genes. We followed the methods described in Sánchez-Cabrera et al. (2021) to map to reference the reads using Geneious v9.0.4 (Kearse et al., 2012). For the outgroups, we used top BLASTn hits (filtering for *Cyclamen* sp., *Camellia* sp., *Vaccinium* sp., *Actinidia* sp. and *Rhododendron* sp. when available) and, when possible, top tBLASTx from *Camellia sinensis*, *Embelia ribes* and *Primula veris* genomes. We reconstructed the relationships using the RAxML 7.2.8 plug-in from within Geneious v9.0.4, searching for the maximum likelihood tree with 1000 bootstrap replicates. We added normalized expression values (TMM) to the phylogenetic trees with “ggtree” (Yu et al., 2017) and “phytools” (Revell, 2012) packages in R v4.0.0 (R Core Team, 2020). We concatenated alignments of all ABP structural and regulatory loci and performed a combined phylogenetic analysis using the same methodology as above but partitioning the data by locus and determining the best fit model of nucleotide substitution.

We calculated the NS SNP rate between blue and orange petals of each species, with a cutoff of 75% difference between colors. We used the isotigs selected for NS SNP search. We focused on genes with orange monophyly to either one or both species, because orange is derived from blue and therefore, we expect genes responsible for the blue to orange shift to show a single common ancestor for the orange samples.

### Differentiating the four hypotheses

To differentiate the four hypotheses (Figure 2) we used the following logic. First, the convergence hypothesis predicts separate origins of the blue to orange color shift which would produce separate clades of orange samples for both species. Under this hypothesis, and with enough data, *La* and *Lm* should be reciprocally-monophyletic. The introgression hypothesis involves a single origin of the color change in both species (after the speciation event) which would appear as very close relationships for the color differentiating locus between orange *Lm* and *La* since they are the same gene, just shared across species boundaries (for the color causing gene(s) the species should not be monophyletic), yet for non-color causing loci, species may be monophyletic. However, the species must be interfertile. Third, the ancestral polymorphism hypothesis produces a single origin of the color change (before the speciation event) creating separate monophyletic blue and orange clades. Finally, if taxonomic confusion has blurred the identity of the species (non-monophyletic species hypothesis, Jiménez-López et al., 2022), then there could be a monophyletic color clade (across species boundaries) – a topology expected for all loci, not just those contributing to the color shift.

### Reproductive isolation between La and Lm

To determine reproductive isolation between *La* and *Lm*, we performed controlled crosses in plants from different populations grown in glasshouse. Flowers were emasculated before anthesis, and hand-pollinations were made during the first day of anthesis within and between species and color morphs. Pollinated flowers were left to set fruits and the fruit production and the number of seeds per fruit were counted.

### Climate niche modeling

To test whether the color morphs of each species differed ecologically across their ranges, we constructed an occurrence dataset using a combination of personal observations and iNaturalist (inaturalist.org) records (manually verified for species identification and color; Appendix 1—figure 6). The dataset comprised 4287 records. All 19 BIOCLIM variables were then extracted for each record based on GPS coordinates at the highest available spatial resolution (30 second) (Fick and Hijmans, 2017). Occurrences with no climate data and duplicated occurrences (multiple occurrences of the same species with the same flower color within a pixel) were deleted. There were 3309 occurrences in the final dataset (*La* blue = 641, *La* orange = 1941, *Lm* blue = 662, *Lm* orange = 65).

To test for univariate climatic differences among the four morphs, a separate Kruskall-Wallis test with a post-hoc Dunn test (non-parametric analogs of ANOVA with post-hoc Tukey’s test) was applied to each variable. Significance tests used Bonferroni adjustment for 76 comparisons (19 climate variables across four species/flower color combinations).

To determine whether similar climate variables were most important for separating color morphs in the two species, logistic regression models were built for each species on a subset on non-correlated climate variables. First, strongly correlated climate variables were removed and a dataset of five variables that were both relatively uncorrelated and putatively biologically meaningful based on field observations and experimental manipulations (Arista et al. 2013; J. Jiménez-Lopez et al., unpublished data) were retained: precipitation of wettest month, precipitation seasonality, isothermality, temperature seasonality and mean temperature of wettest quarter. Data were normalized using the ‘normalize’ procedure in the ‘decostand’ function from the vegan R package (Oksanen et al. 2019) to facilitate later comparison of regression coefficients. Best fit models were selected using AIC scores derived from the ‘dredge’ function from the MuMIn R package (Bartón, 2020).

To test whether the two orange morphs were more divergent than the two blue morphs across the 19 BIOCLIM variables, a randomization procedure was used. The specific null hypothesis was no difference between the distance between the two orange morph means and the distance between the two blue morph means. To build a null model for each of the 19 variables, species identities were retained but color morphs were randomized 999 times. For each distribution, one tail indicated that differences between orange populations was greater while the other tail indicated greater differences between blue populations. A two-tailed test indicated whether observed orange differences were greater than expected, whether blue differences were greater than expected or whether neither color was more divergent.

## Data Availability

The data that support the findings of this study are openly available in Genbank (raw reads & assembled contigs) and Figshare (spectra, biochemistry, expression values, alignments). Raw reads, assembled contigs and expression values will be made available from Genbank and FigShare. R scripts to conduct statistical analyses will also be available from FigShare.

## Acknowledgements

We would like to thank the support staff in the Department of Biology and the Department of Chemistry and Biochemistry at Santa Clara University. Boris Igić provided thoughtful insights on the evolutionary history of color in these species. We would like to thank M. Abdelaziz for his help in organizing the field collecting trip and during the sampling of Moroccan *L. monelli* plants. Grants PID2020-116222GB-100 to M.A. and E.N., CGL2015-63827 to M.A. and P.L.O. and BES-C-2016-0023 to M.S.C funded by MCIN/AEI/ 10.13039/501100011033 and by “European Union NextGenerationEU/PRTR”.

## Author contributions

MS-C, EN, MA, PO, and JW designed and performed research; AF conducted the UHPLC technique; BC conducted the climate niche analyses; FJ-L performed plant crosses and supplied *L. arvensis* reflectance data. MS-C, EN, and JW analyzed data; and MS-C and JW wrote and revised the paper with assistance from coauthors.

## Appendix 1

**Appendix 1—figure 1.**
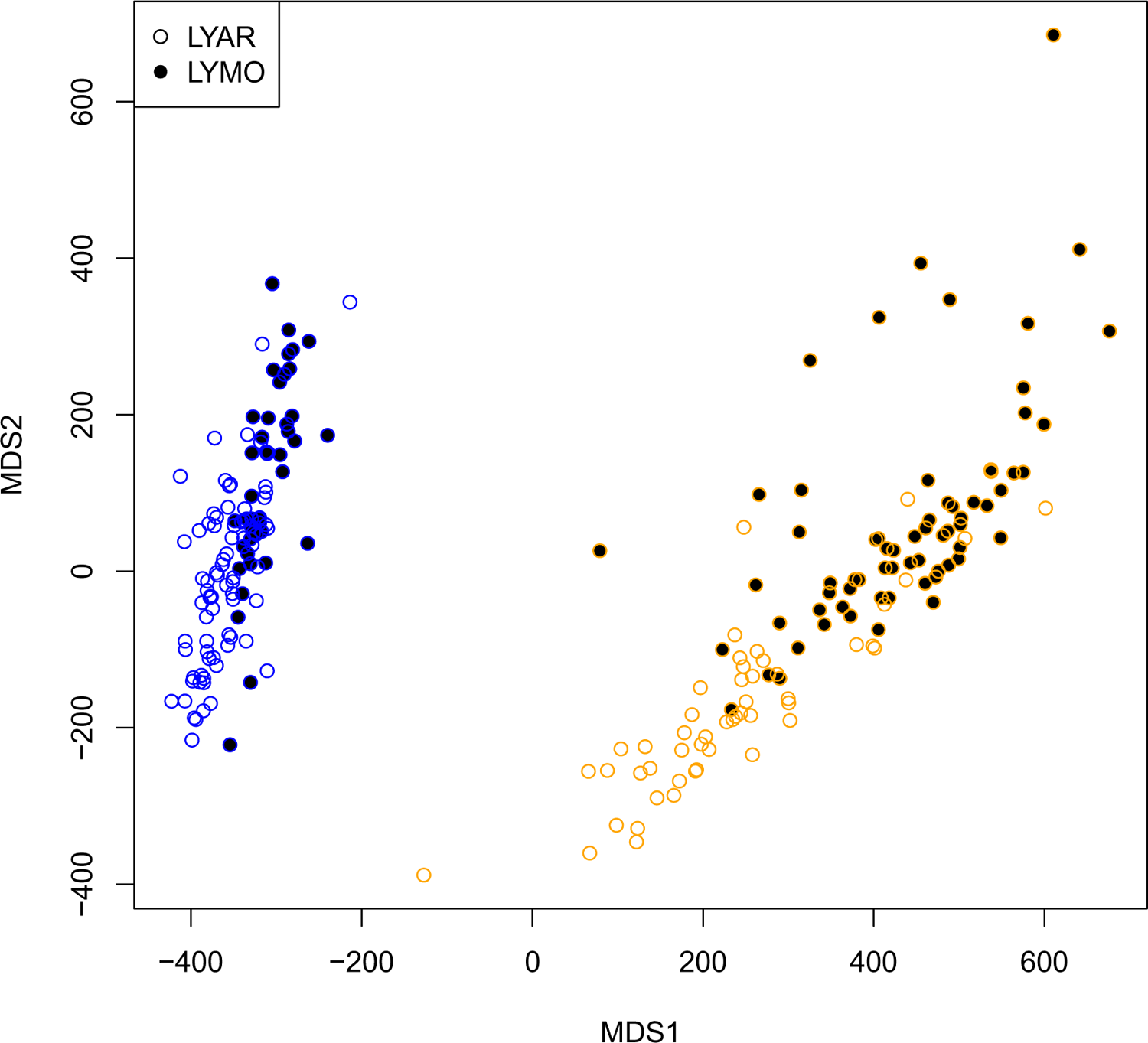
NMDS plot of UV-Vis reflectance spectra for blue and orange petalled individuals of *L. arvensis* (LYAR) and *L. monelli*. Statistical analysis was assessed with permutational manova (Anderson, 2001) with the following results (1) restricting permutations within color (comparing across species), R^2^ = 0.15113, F = 41.662, p < 1e-04 vs. (2) restricting permutations within species (comparing across colors), R^2^ = 0.74999, F = 701.98, p < 1e-04. Very similar results were found after applying a brightness correction – subtracting the minimum reflectance value of each sample from all wavelengths (not shown).

**Appendix 1—figure 2.**
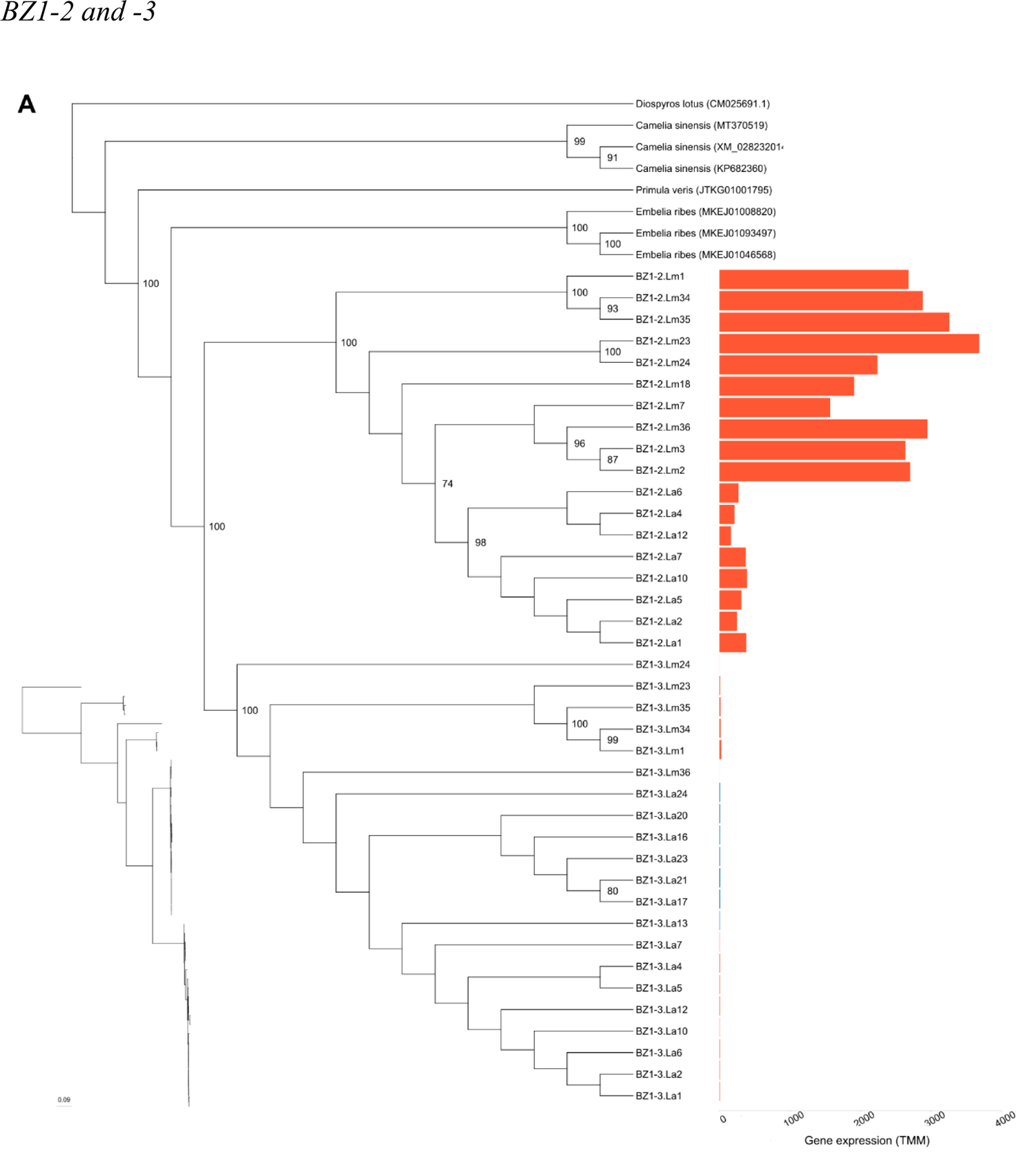
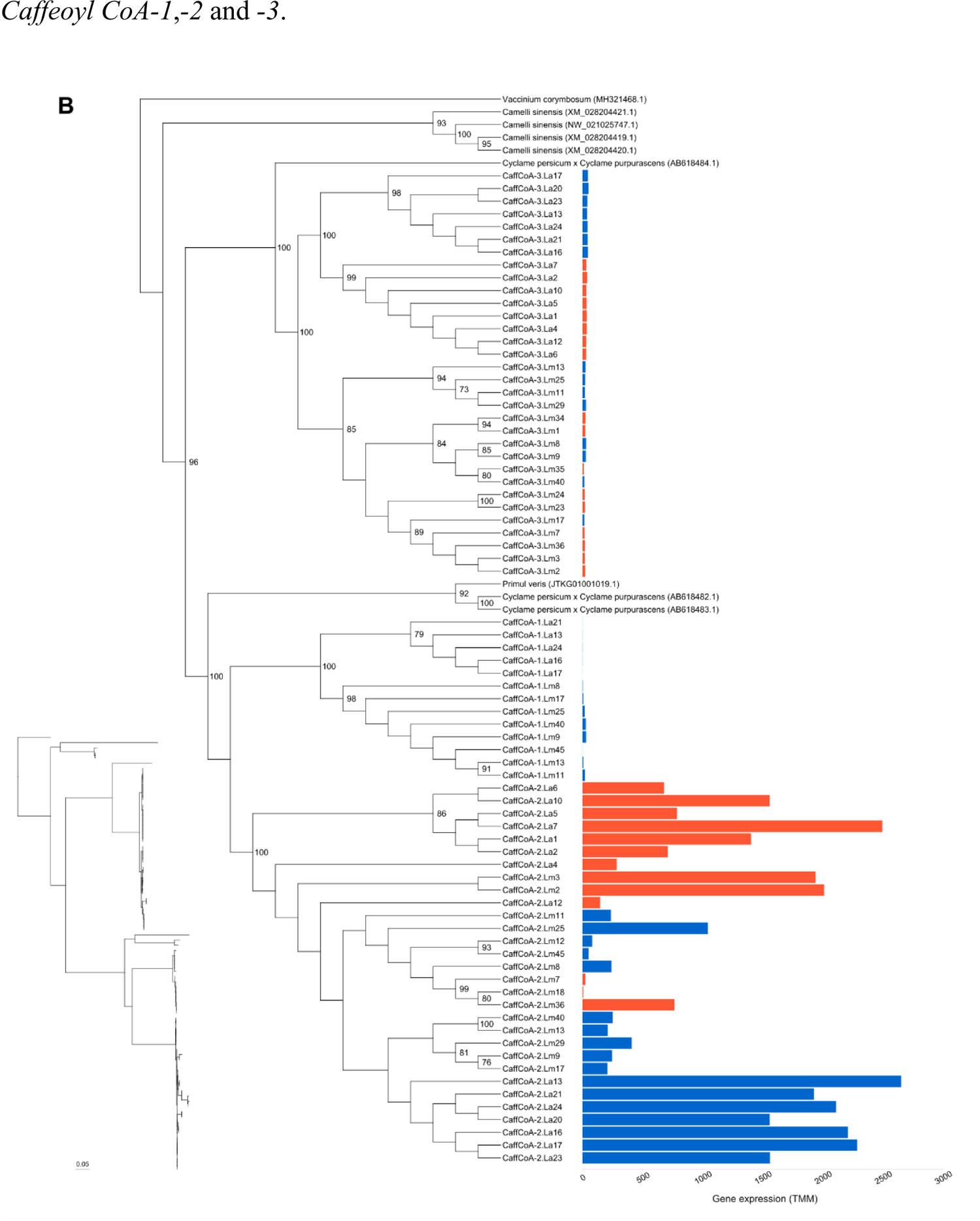
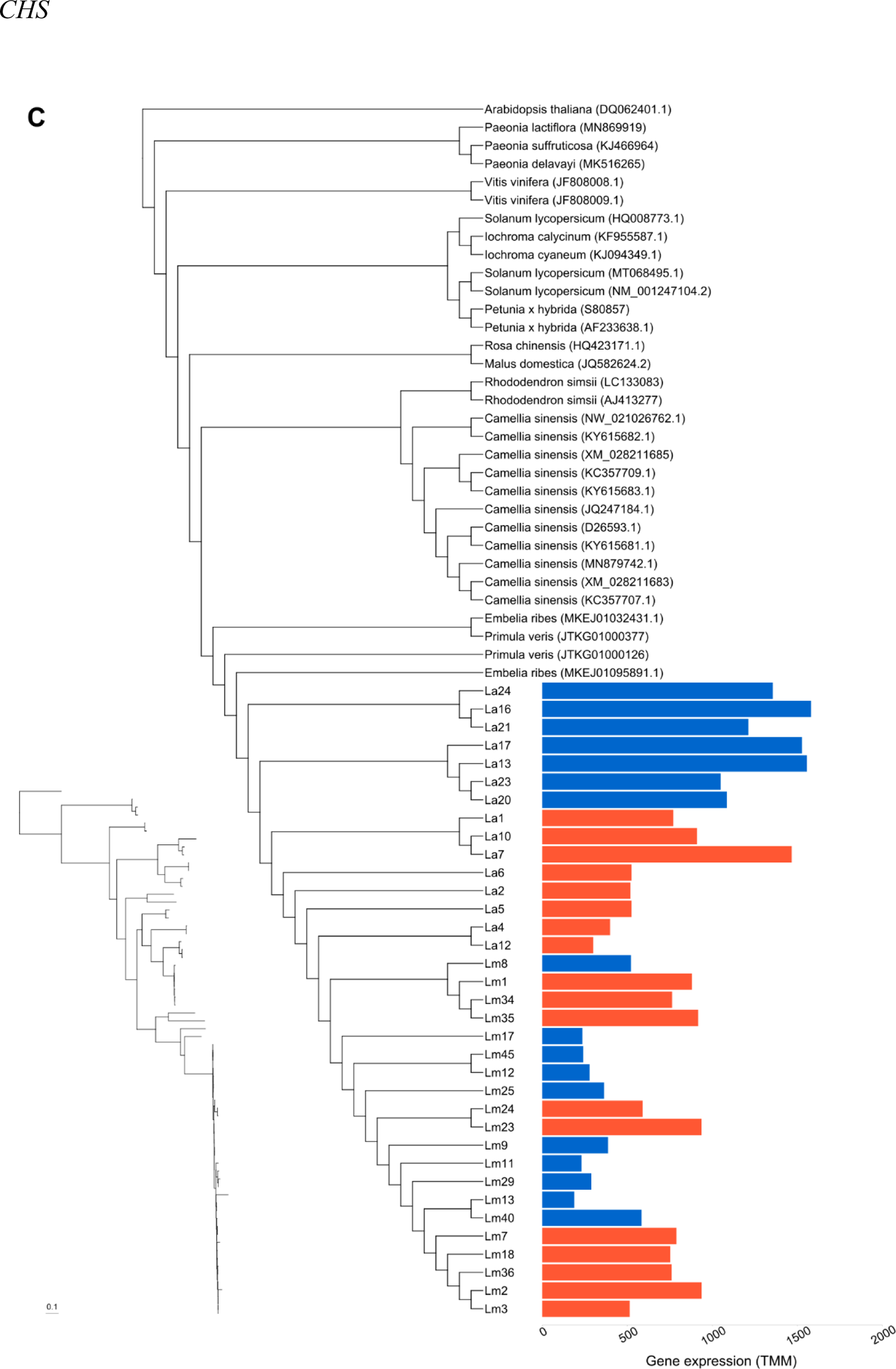
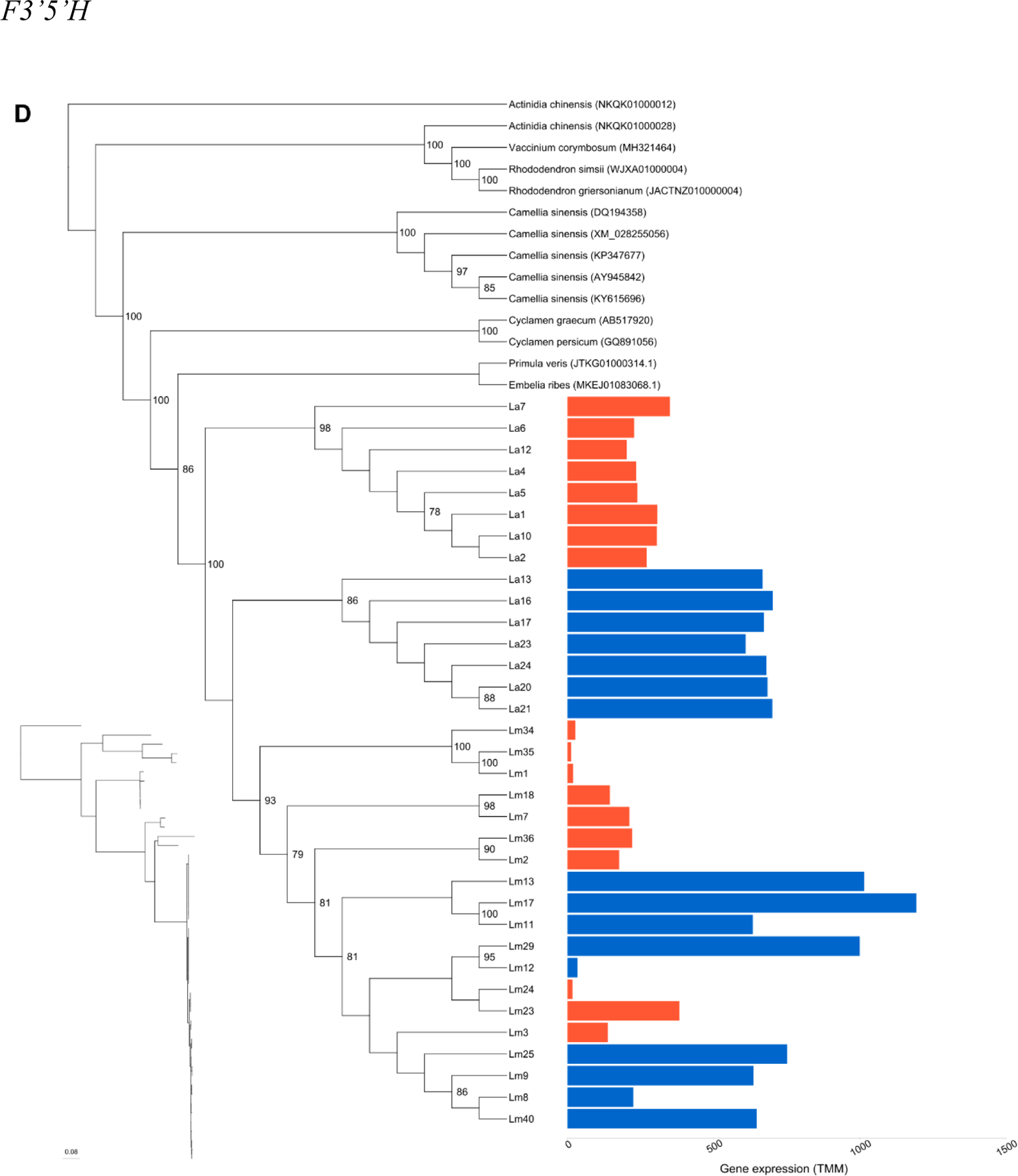
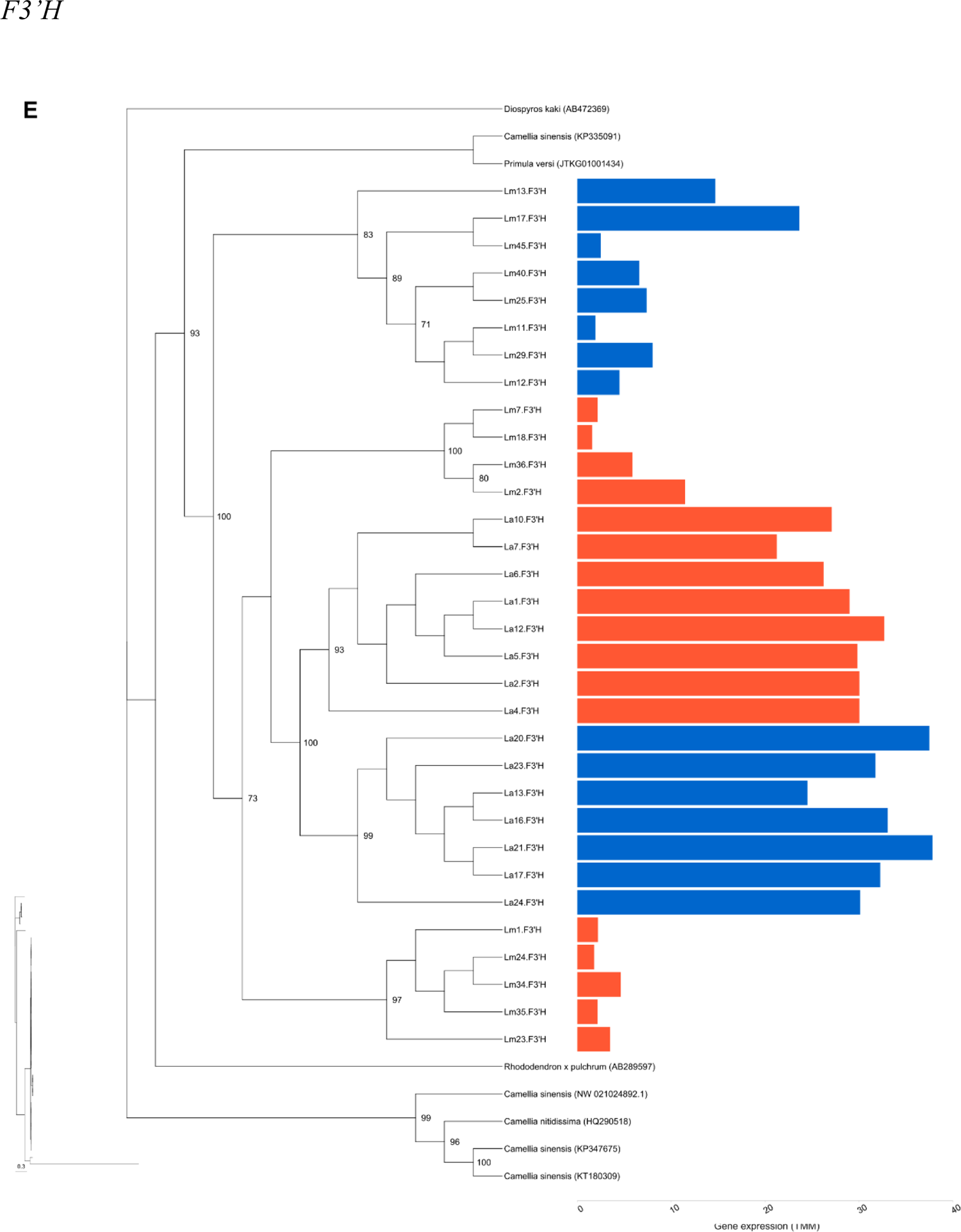
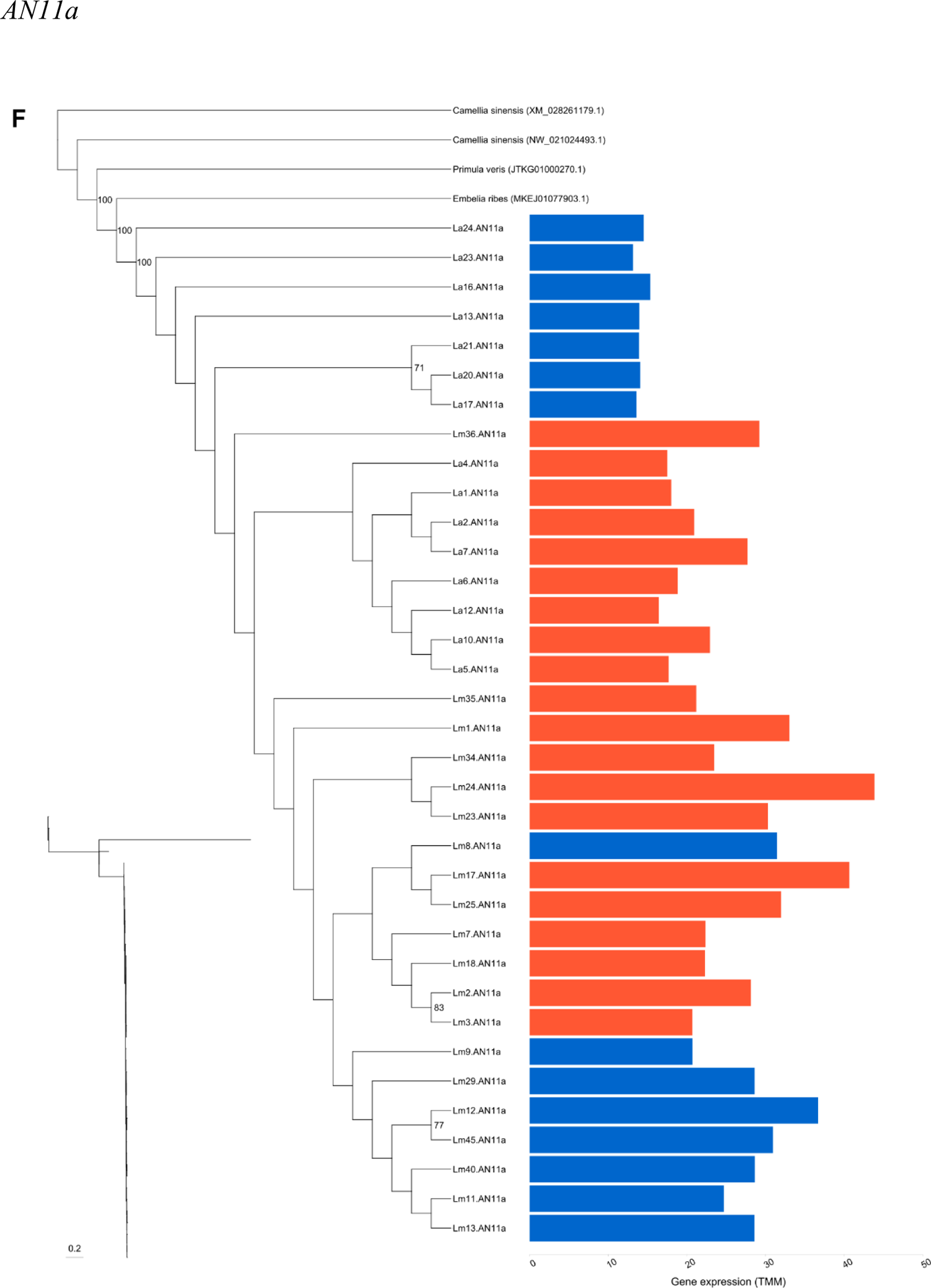
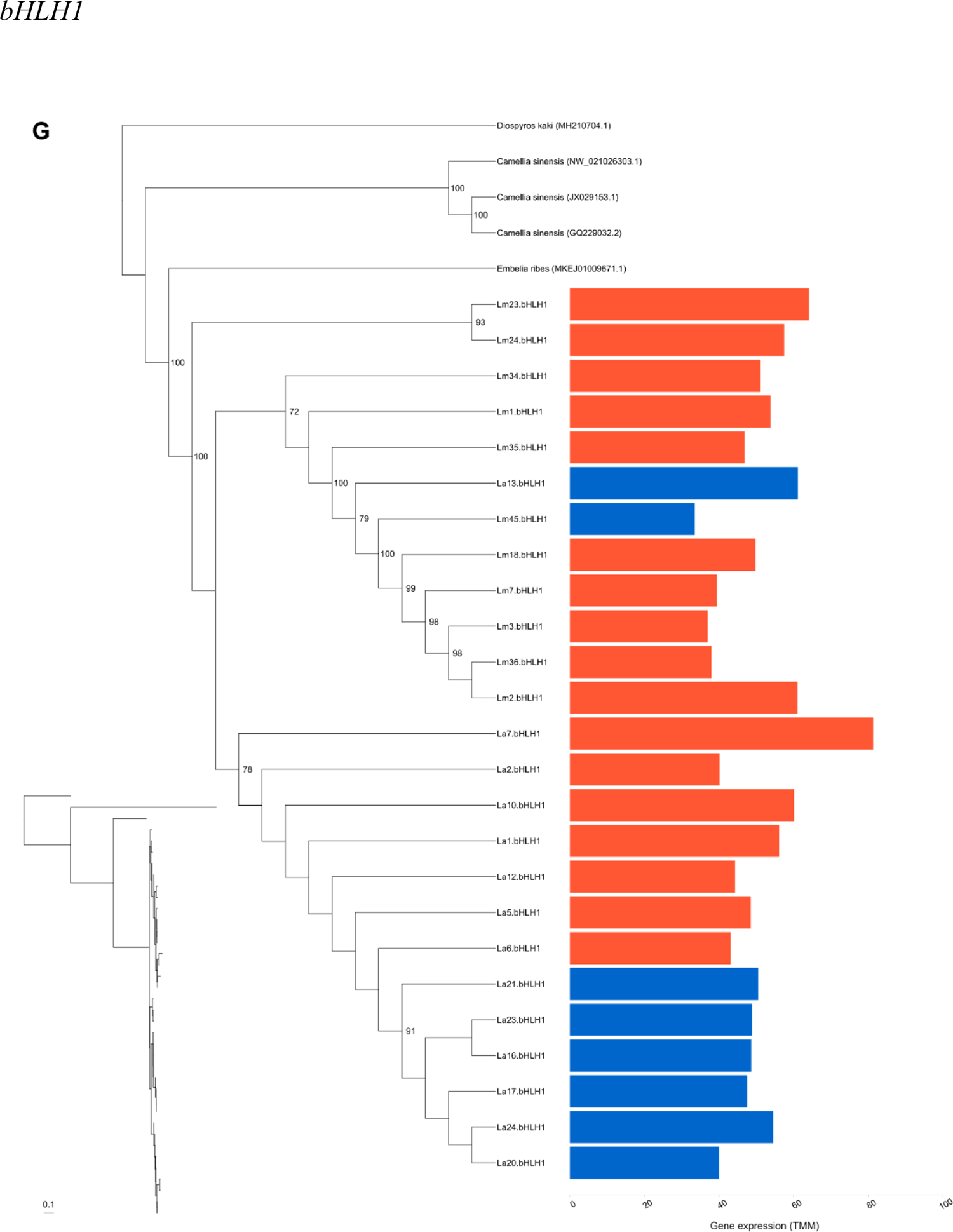
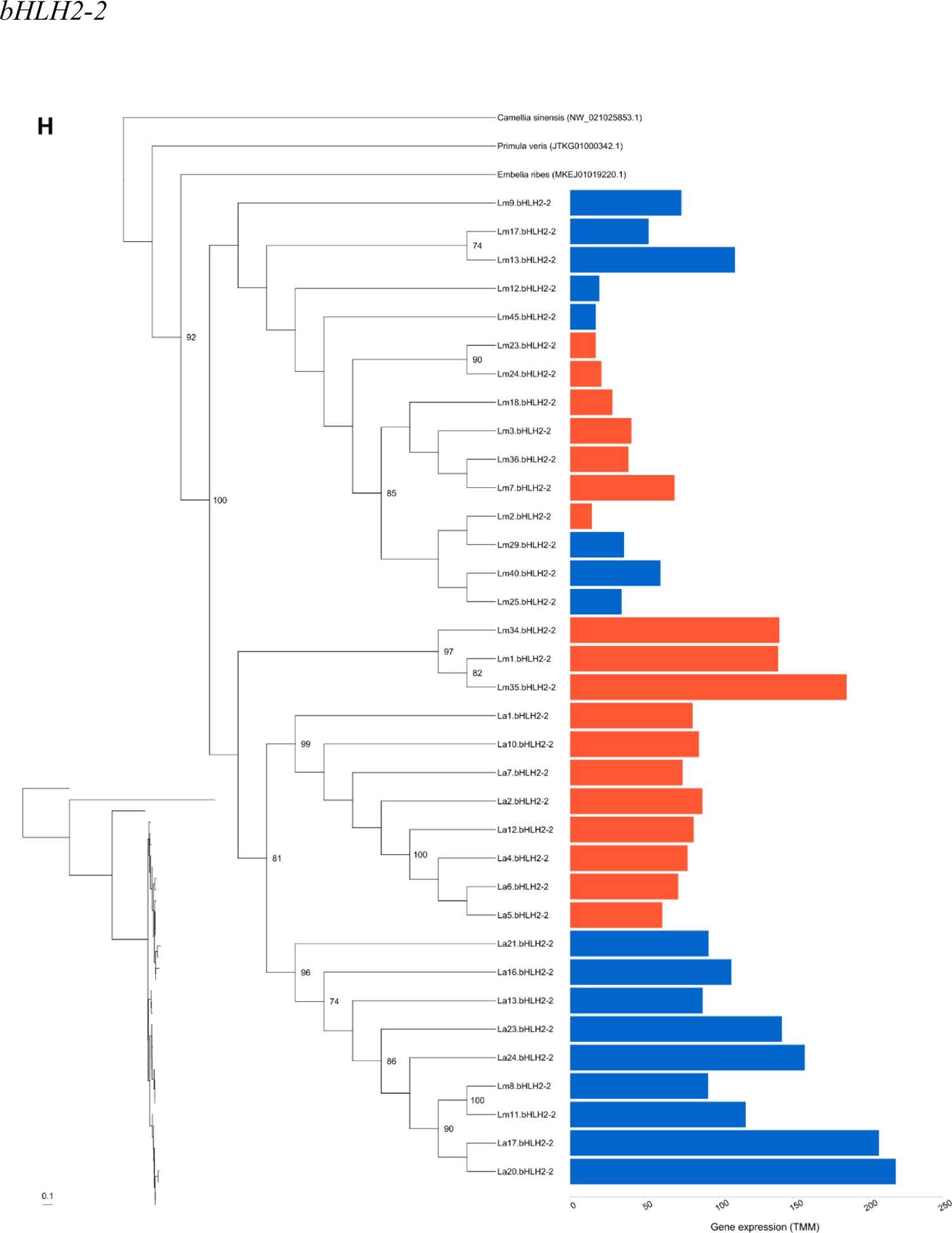
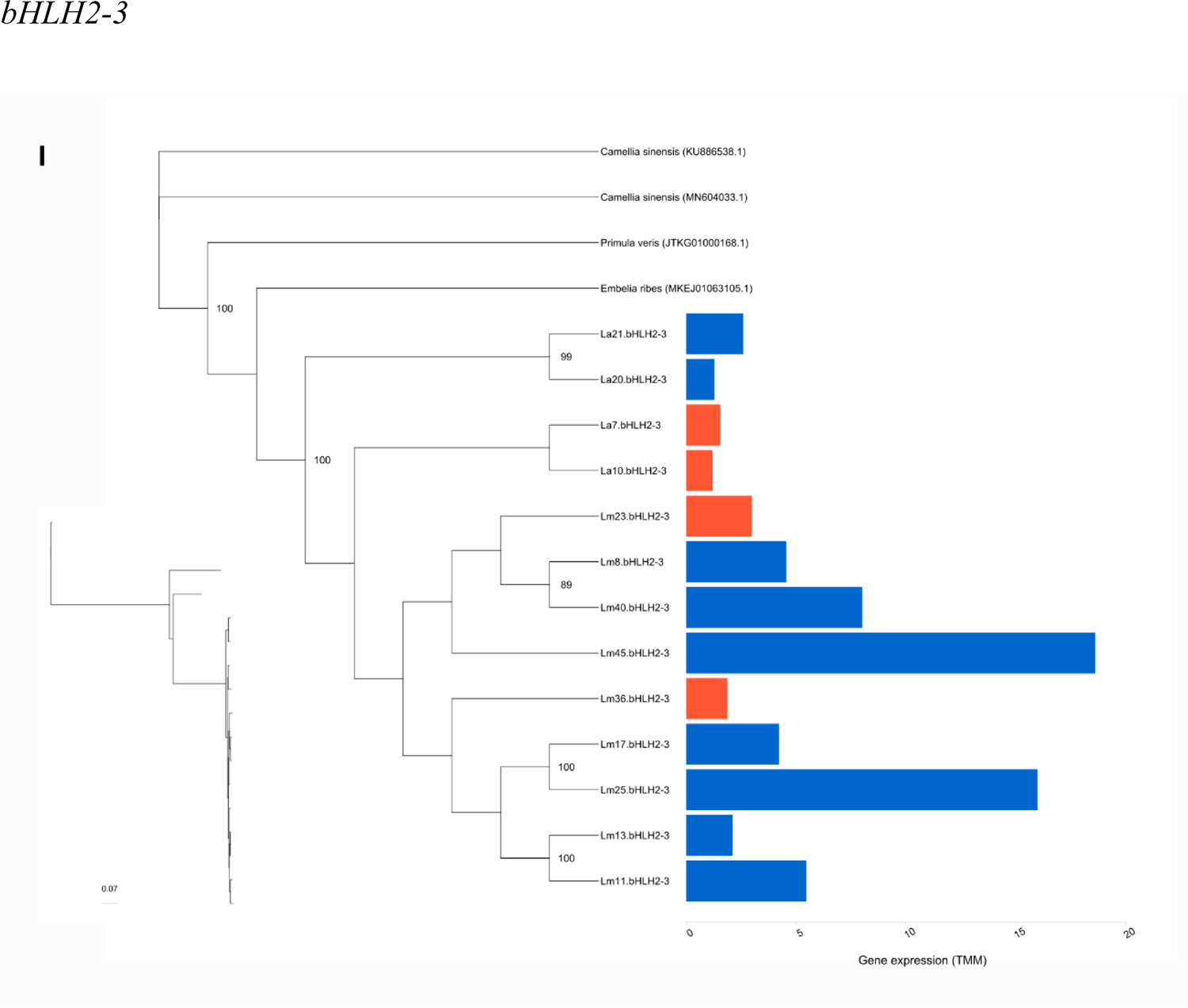
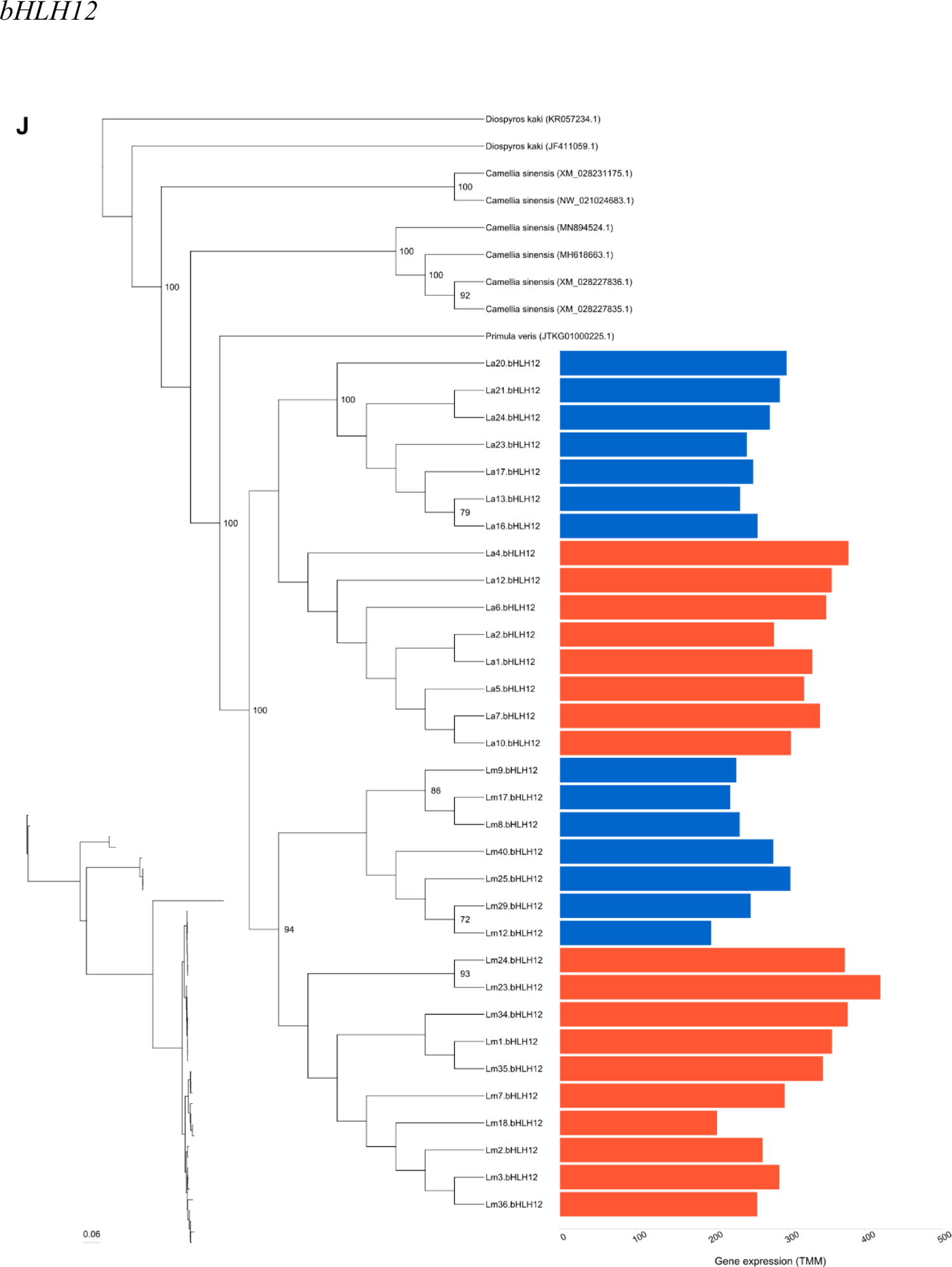
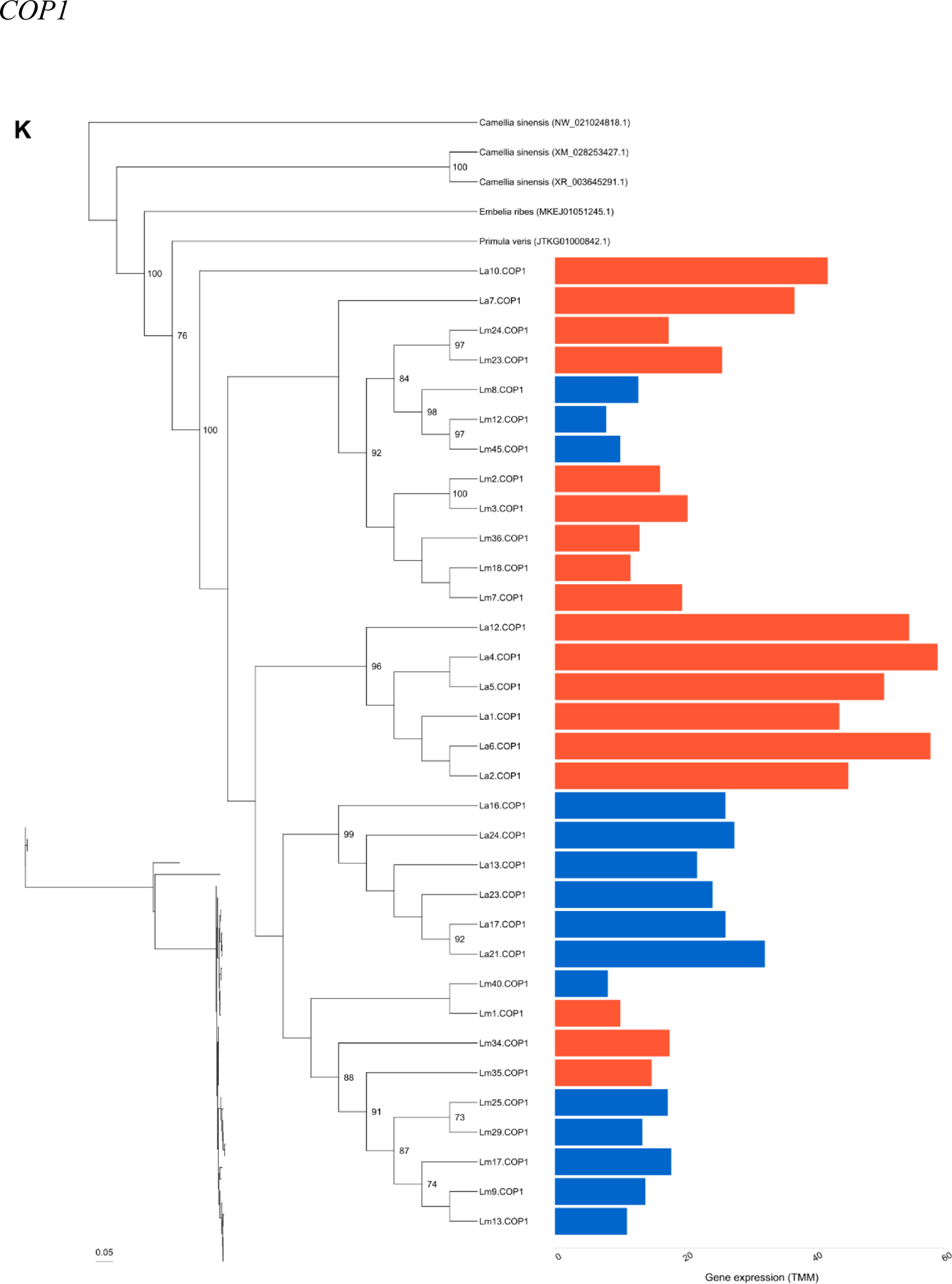
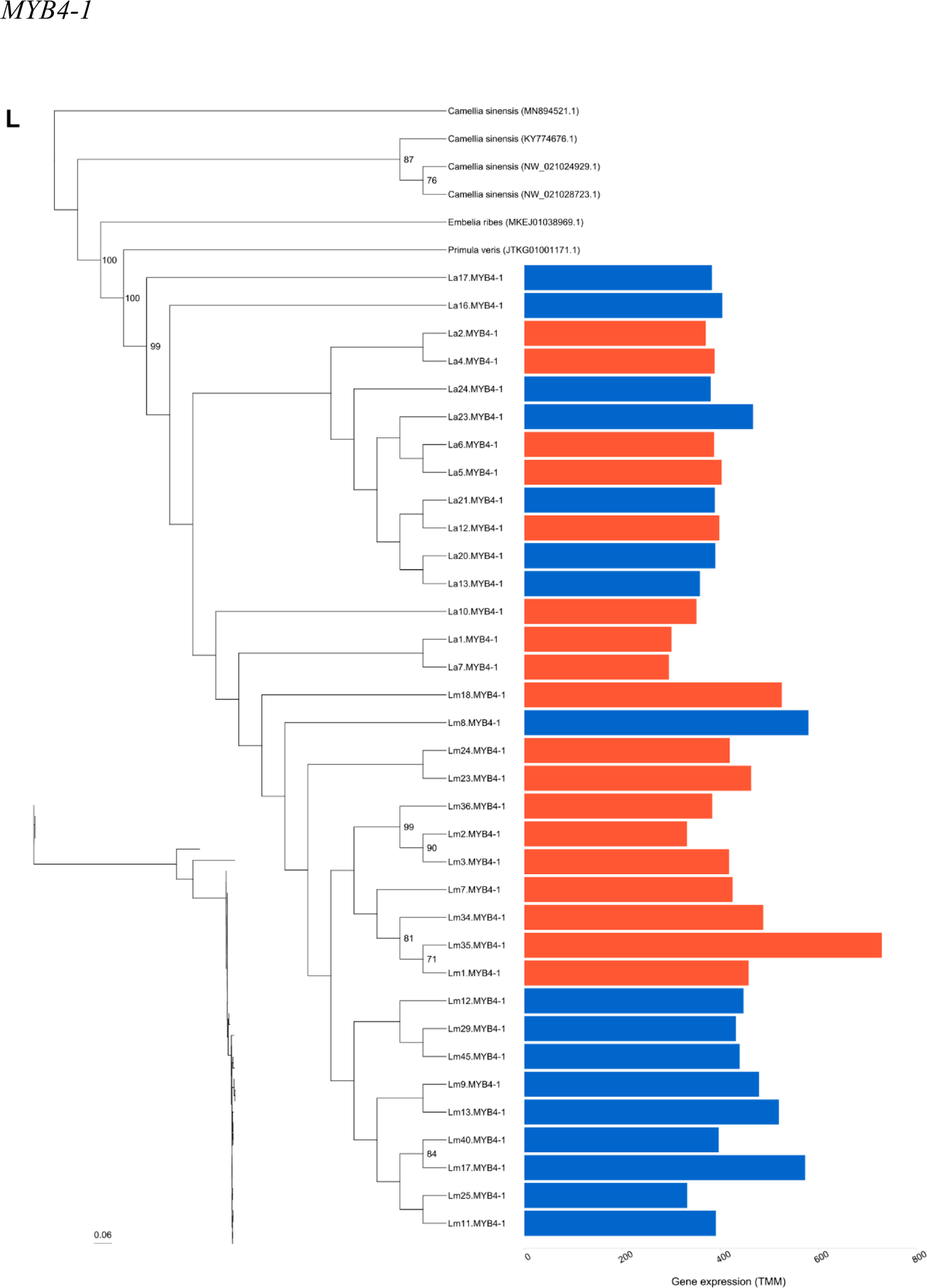
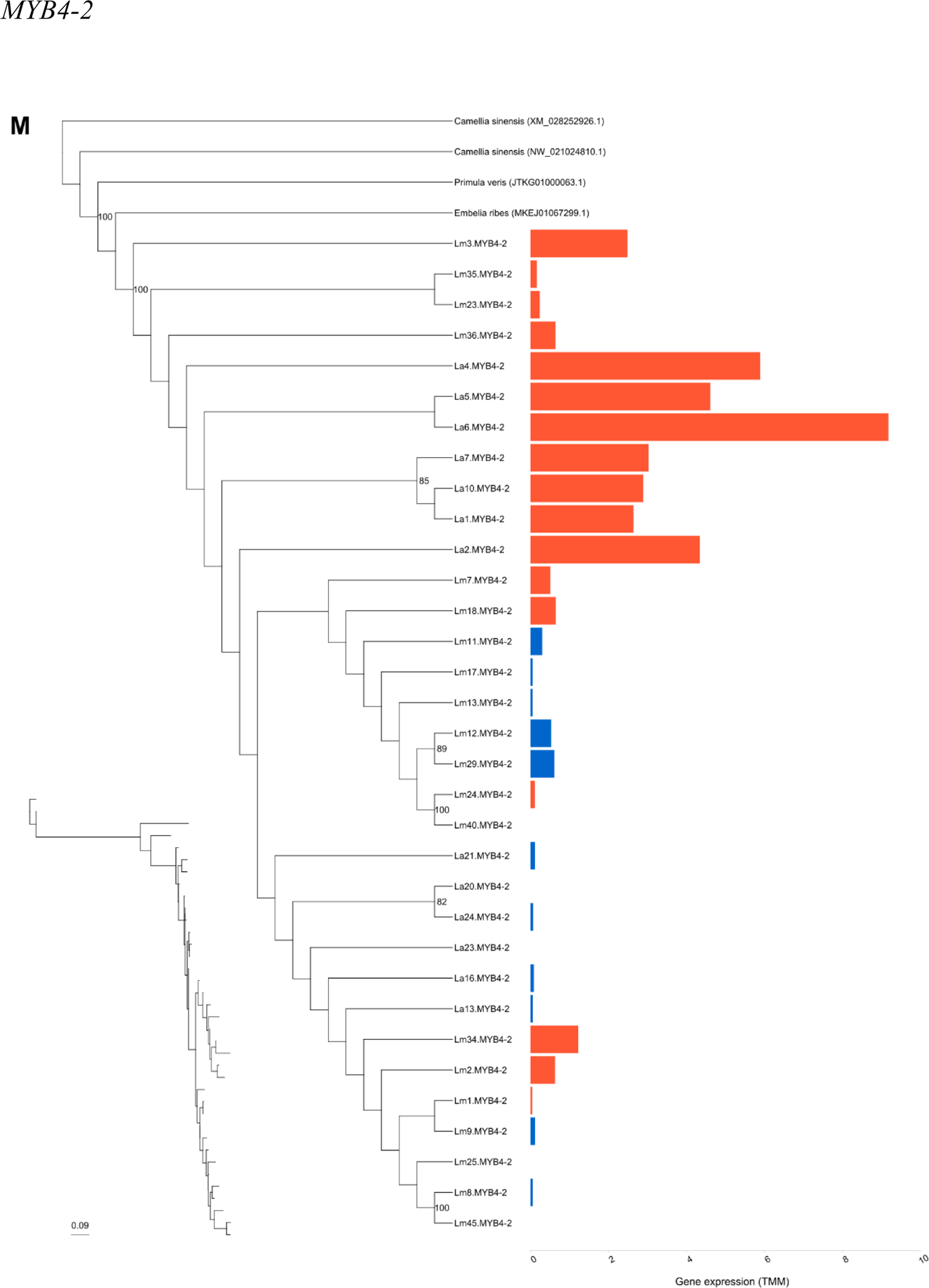
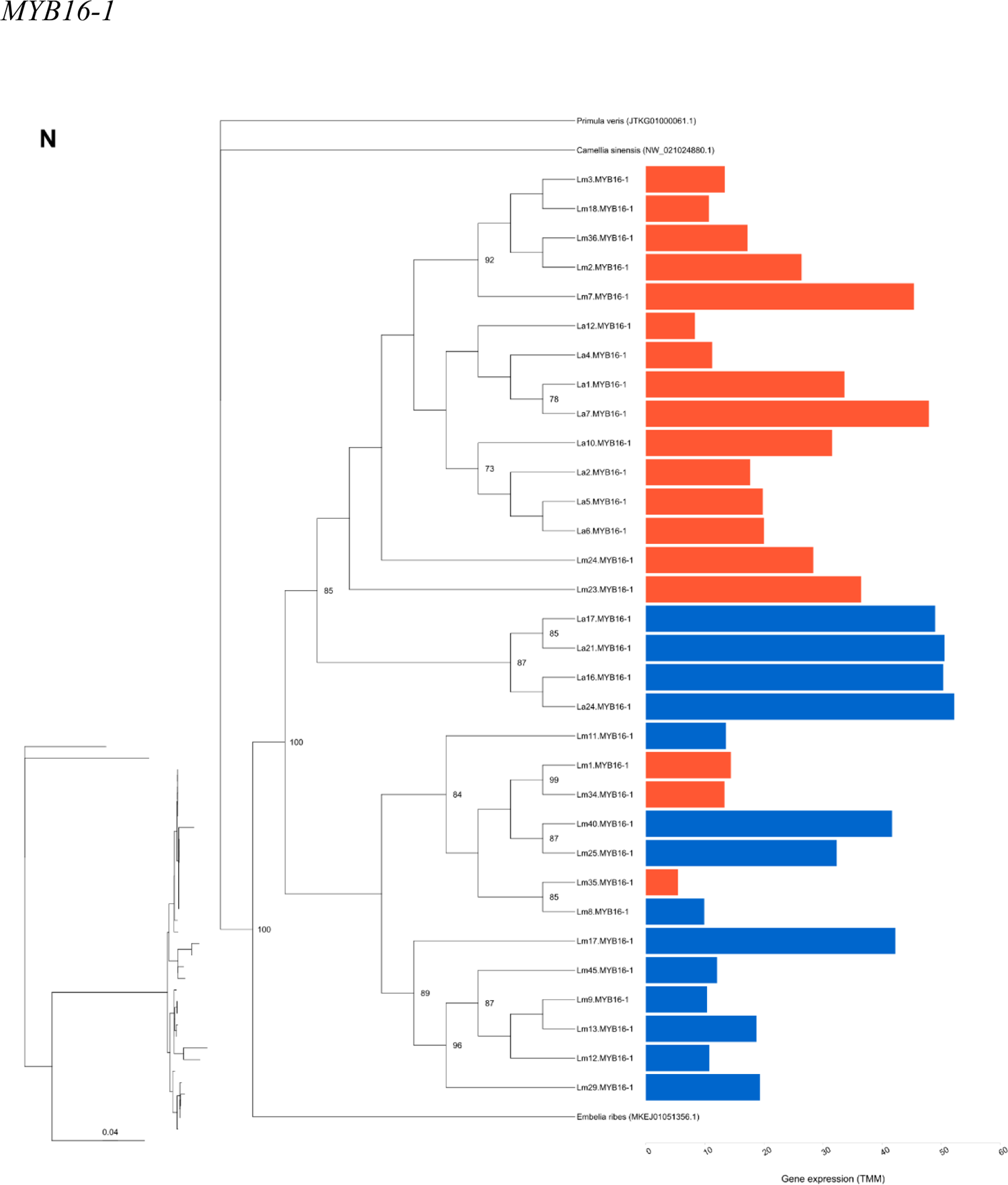
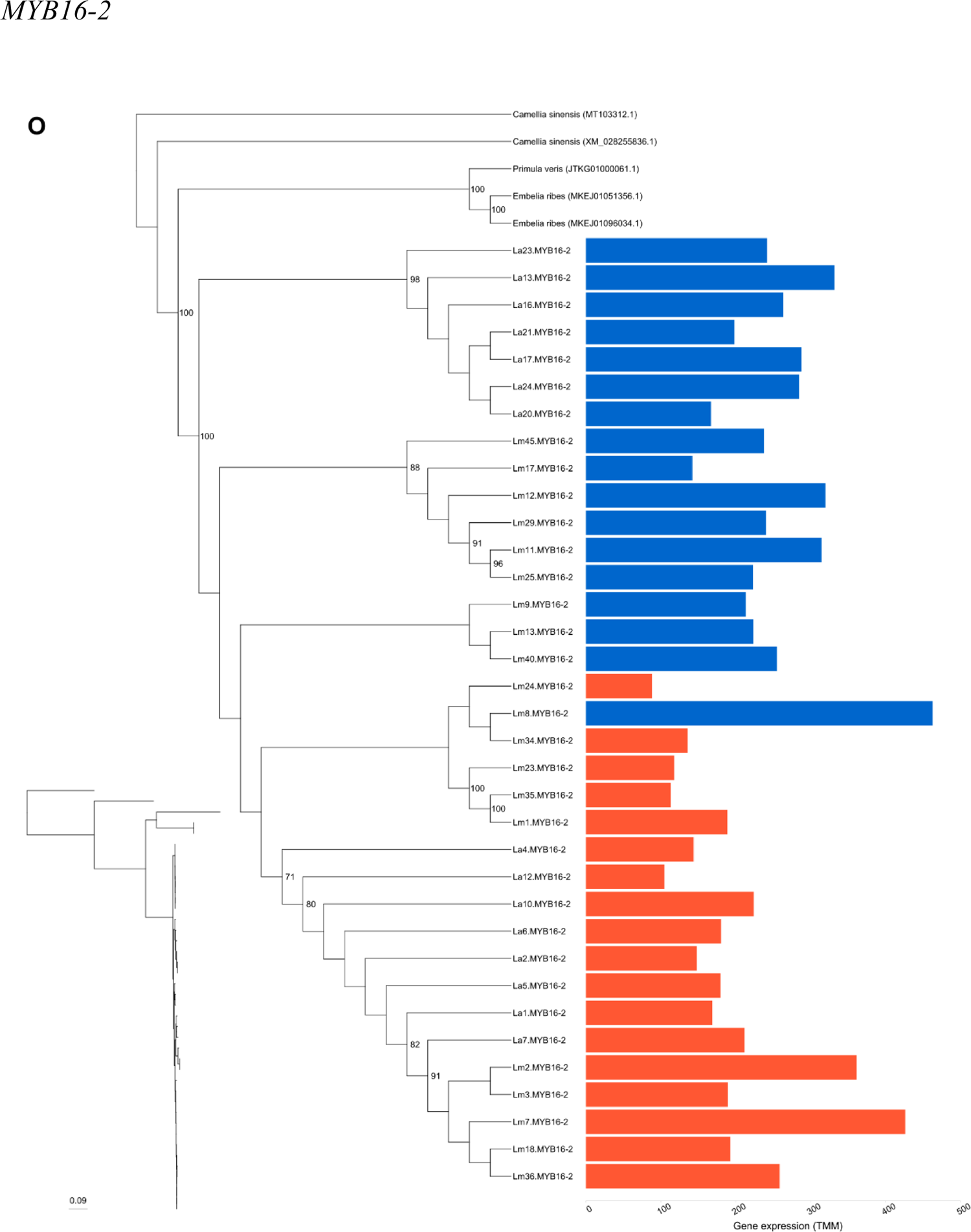
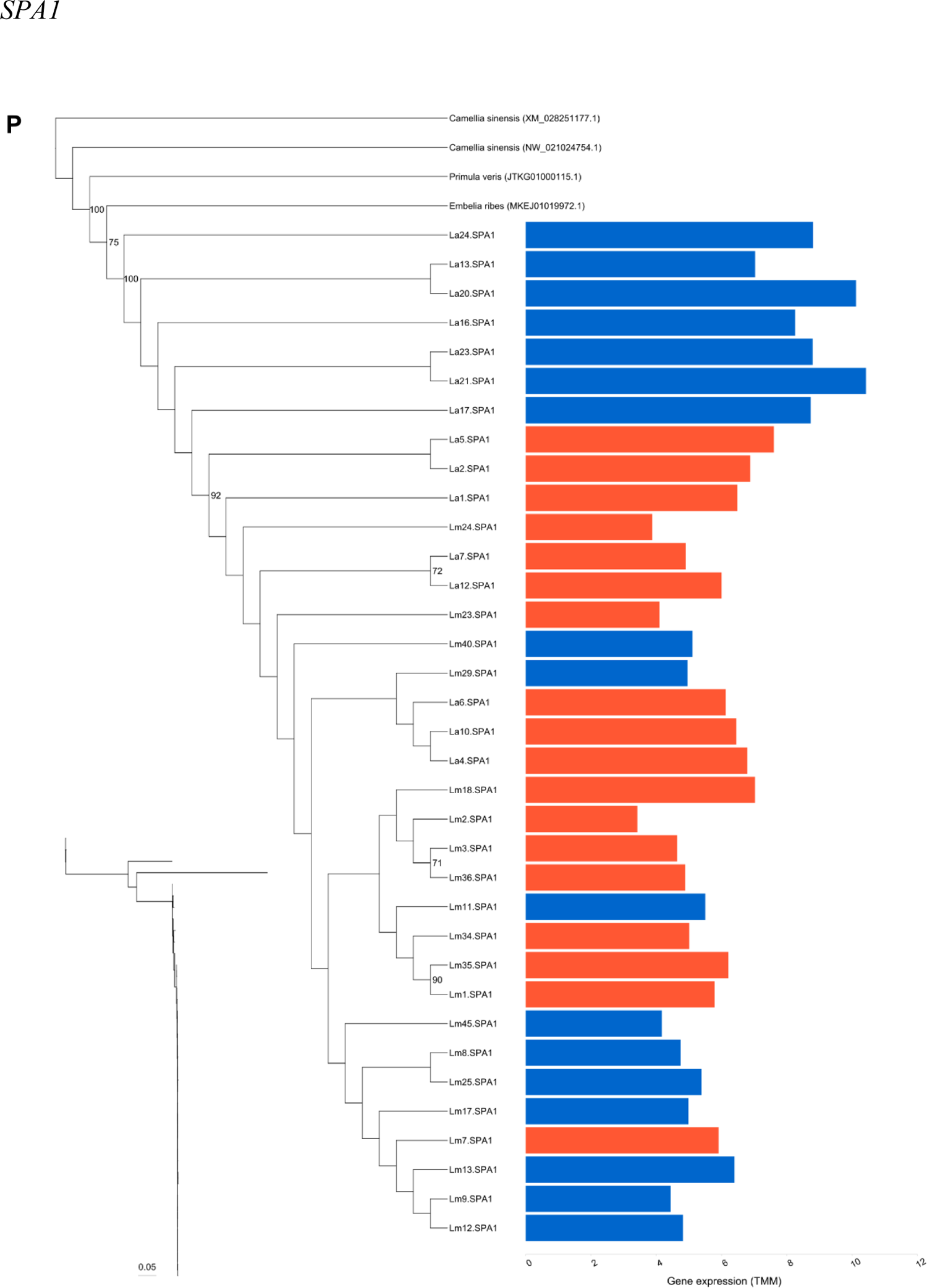
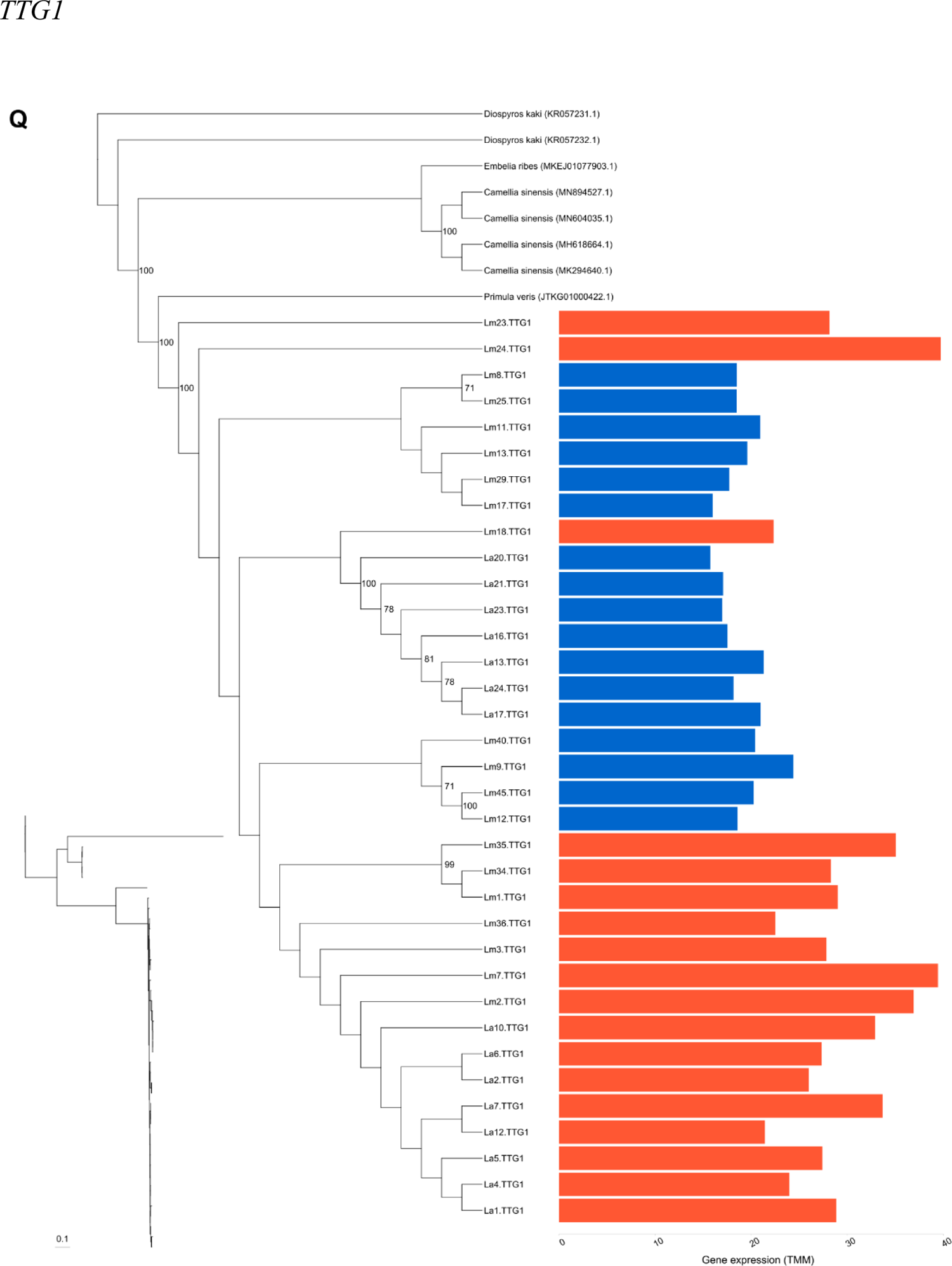
Maximum likelihood phylogenetic analysis of the coding sequence for eight structural (A-E) and 12 regulatory (F-Q) loci from blue and orange petals of *L. arvensis* and *L. monelli*. (A) *BZ1-2* and *-3*. (B) *Caffeoyl CoA-1*,*-2* and *-3*. (C) *CHS*. (D) *F3’5’H*. (E) *F3’H*. (F) *AN11a*. (G) *bHLH1*. (H) *bHLH2-2*. (I) *bHLH2-3*. (J) *bHLH12*. (K) *COP1*. (L) *MYB4-1*. (M) *MYB4-2*. (N) *MYB16-1*. (O) *MYB16-2*. (P) *SPA1*. (Q) *TTG1*. Outgroups were included when available from the genomes of closely related species (*Camelia sinensis, Embelia ribes* and *Primula veris*). Bootstrap values above 70% are provided to the right of the nodes. Inset phylograms are provided for branch length comparisons with samples in same order as the larger cladogram and contain a scale bar in substitutions per site. The bar plots show the gene expression level (TMM values) of blue and orange flowers.

**Appendix 1—figure 3.**
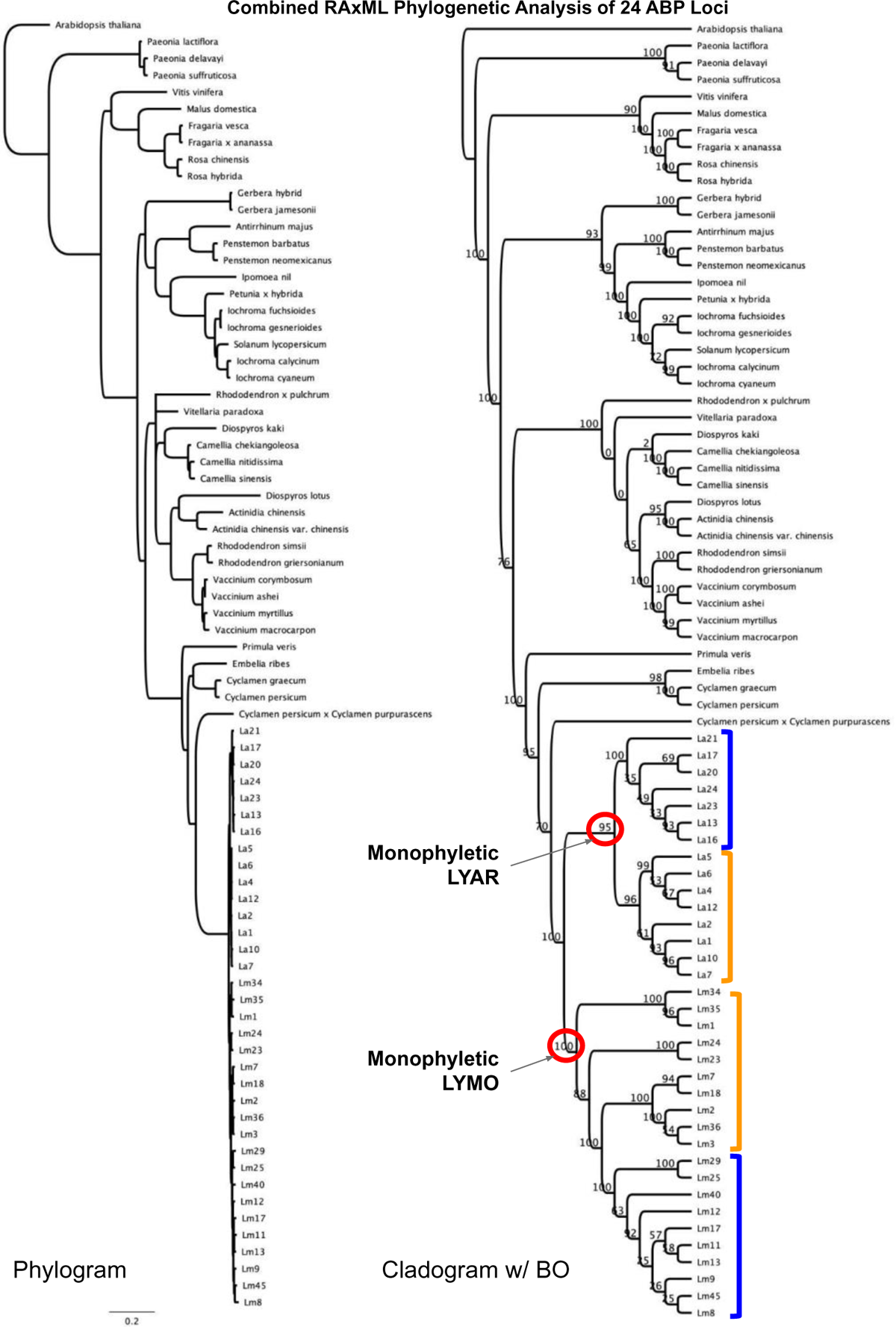
Combined maximum likelihood phylogenetic analysis of 24 ABP structural and regulatory loci. Phylogram on the left with branchlengths and cladogram on the right with bootstrap values indicated at the nodes. Cladogram annotated with petal color (blue and orange brackets). Lysimachia arvensis (LYAR and La samples) and L. monelli (LYMO and Lm samples) are strongly supported as monophyletic as are the color types within each species except for the paraphyly of LYMO orange.

**Appendix 1—figure 4.**
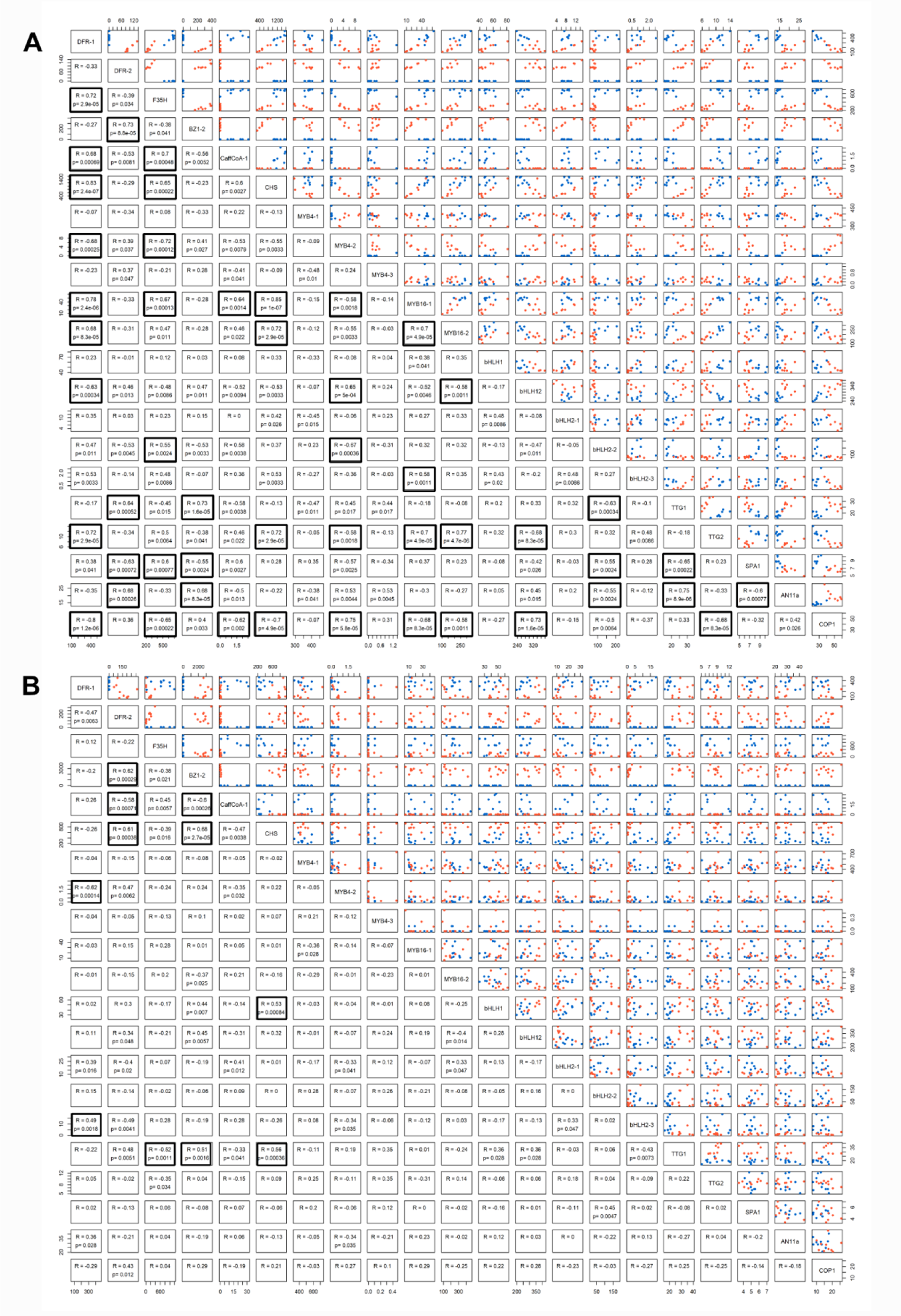
Kendall correlations of the expression level (TMM values) between the most relevant differential expressed structural genes (*DFR-1*, *DFR-2*, *F3’5’H*, *BZ1-2*, *CaffCoA-1*, *CHS*) and the regulatory genes (*MYB4-1*, *MYB4-2*, *MYB16-1*, *MYB16-2*, *bHLH1*, *bHLH12*, *bHLH2-1*, *bHLH2-2*, *bHLH2-3*, *TTG1*, *TTG2*, *SPA1*, *AN11a*, *COP1*) present in blue and orange *L. arvensis* (A) and *L. monelli* (B). Correlation coefficients (R) and P values are listed below the diagonal and the raw data above the diagonal. Black boxes indicate those correlations that are significant following a Bonferroni correction.

**Appendix 1—figure 5.**
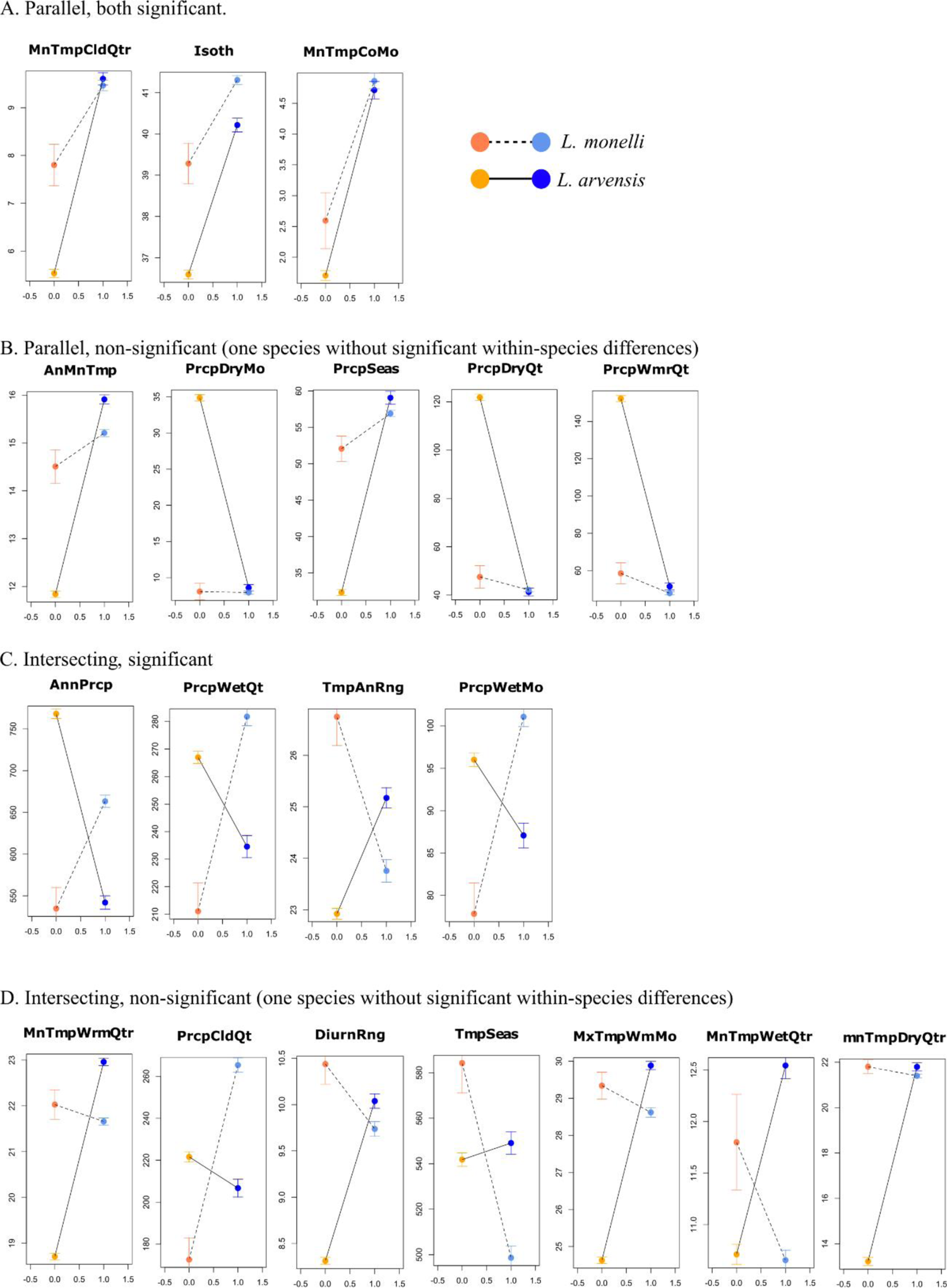
Climate niche modeling results for blue and orange morphs of *L. arvensis* and *L. monelli.* Grouped the 19 variables into 4 categories. (A) Parallel significant: slopes are similar (both + or both -), with orange/blue *L. monelli* significantly different AND orange/blue *L. arvensis* significantly different. (B) Parallel non-significant: slopes are similar, one or both within-species comparison is non-significant. (C) Intersecting significant: slopes are different (one +, one -), within-species differences are significant. (D) Intersecting non-significant: Slopes are different, one within-species comparisons is non-significant.

**Appendix 1—figure 6.**
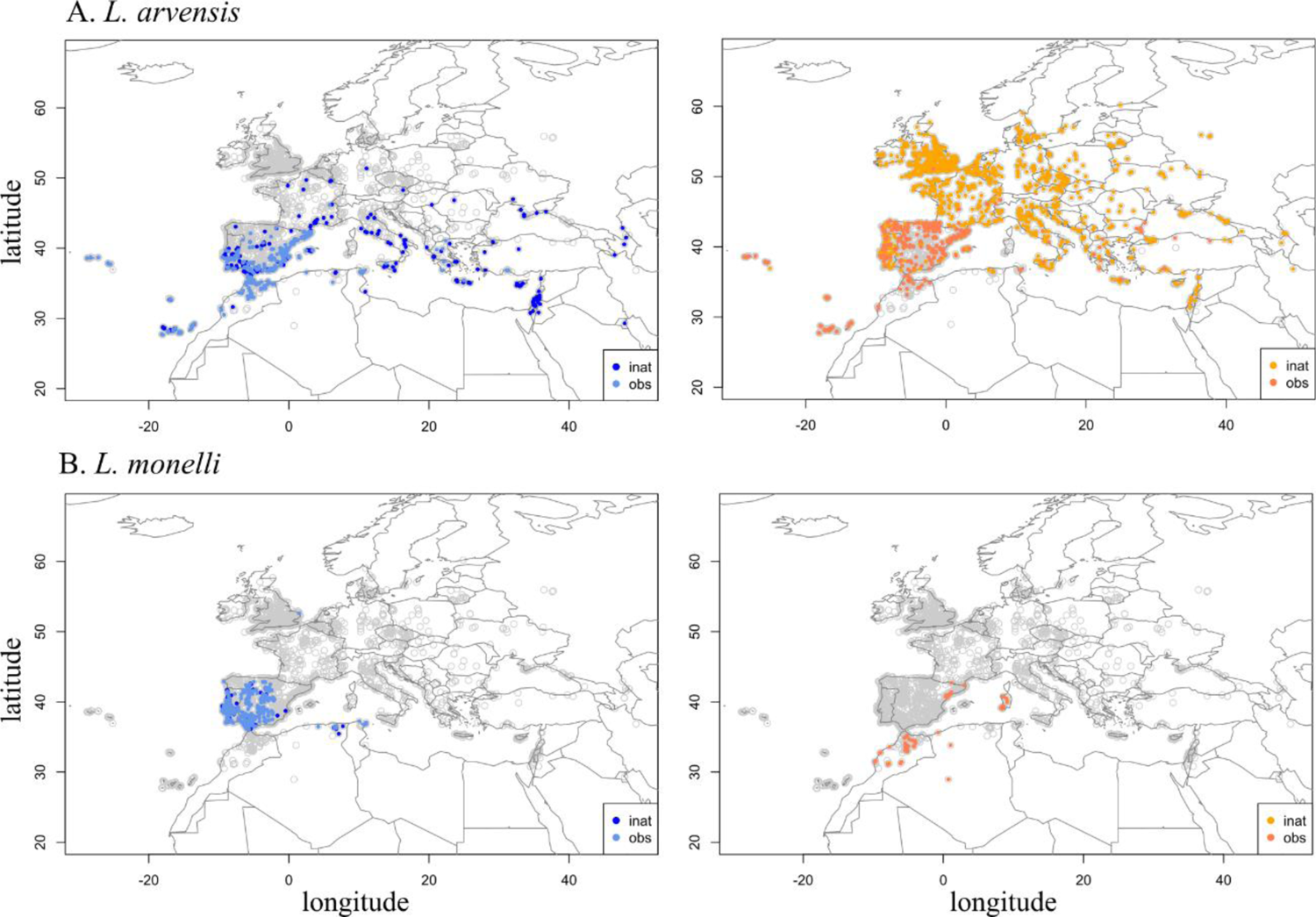
Occurrence data used in climate niche model from individually reviewed iNaturalist records and other sources (herbarium records, personal observations). For *L. arvensis*, there are 1941 orange occurrences and 641 blue (A). For *L. monelli*, there are 662 blues and 65 oranges.

---

**Appendix 1—table 1.**
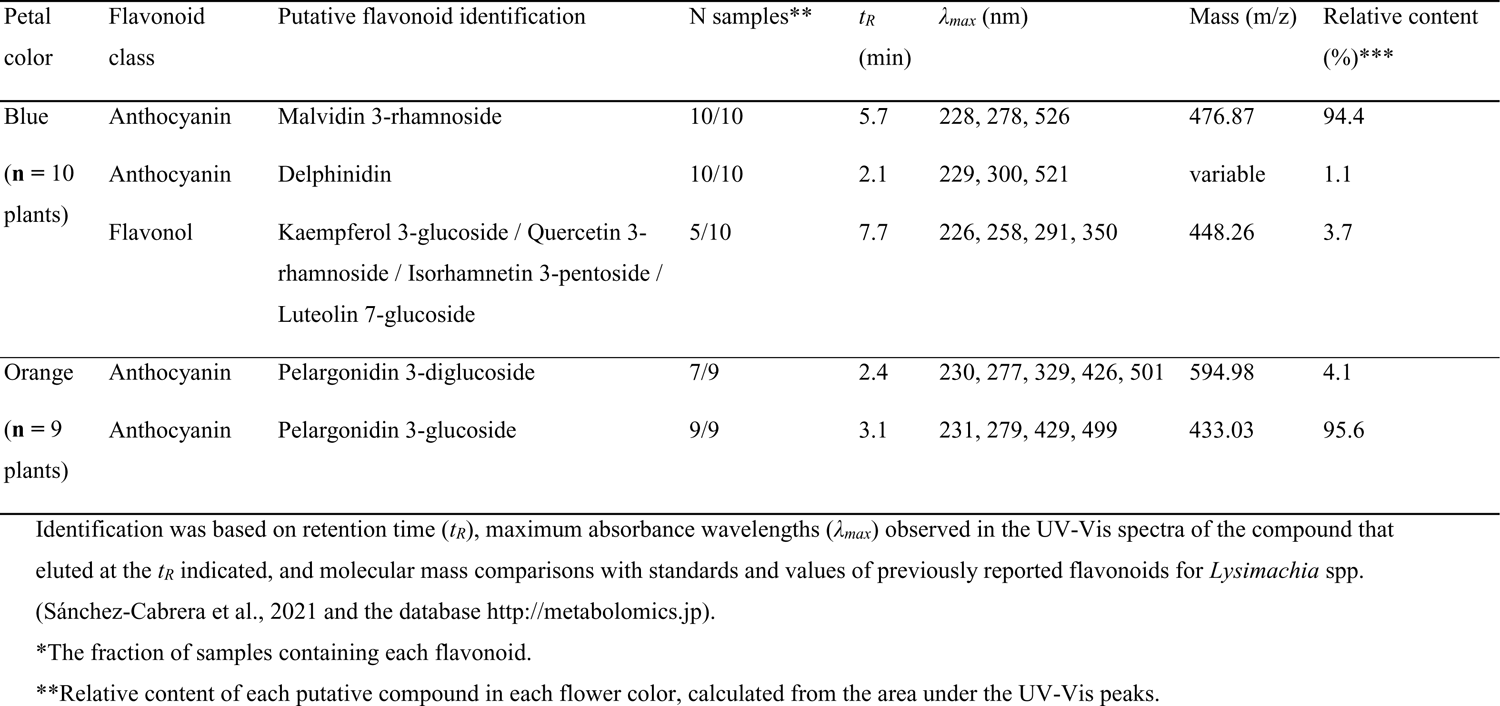
Putative flavonoid identifications of blue and orange petal extracts of *L. monelli* from the UHPLC–MS biochemical analysis (MS analysis was acquired in positive mode). Flavonoids <1% (trace) are not shown.

**Appendix 1—table 2.** Flavonoid biosynthetic pathway genes with significant differential expression between blue- and orange-flowered *L. monelli*.

| Gene<br>abbreviation | Gene name | Blue<br>expression<br>(TMM) | Orange<br>expression<br>(TMM) | Log fold-change | Pathway |
| --- | --- | --- | --- | --- | --- |
| *BZ1-2* | *Anthocyanin 3-O-glucosyltransferase* | 0.31 | 2651.84 | -13.05 | ABP |
| *DFR-2* | *Dihydroflavonol 4 reductase* | 0.01 | 157.18 | -12.53 | ABP |
| *CHS* | *Chalcone synthase* | 331.90 | 784.13 | -1.24 | ABP |
| *Beta-glucosidase* | *Beta-glucosidase* | 0.09 | 4.13 | -5.37 | Non-ABP |
| *Caffeoyl CoA-1* | *Caffeoyl CoA-O-methyltransferase* | 12.73 | 0.32 | 5.29 | Non-ABP |

**Appendix 1—table 3.** Summary of individual phylogenetic analyses of ABP structural and regulatory genes with high bootstrap support (>70%) indicating monophyletic clades in specific groups (genus, species, color). Genes showing orange monophyly (either for *L. arvensis*, *L. monelli* or both species) are highlighted in bold.

| Genes | *Lysimachia* | La | La-O | La-B | Lm | Lm-O | Lm-B | O | B |
| --- | --- | --- | --- | --- | --- | --- | --- | --- | --- |
| Structural genes |  |  |  |  |  |  |  |  |  |
| *BZ1-2\*\** | 100 | 98 | - | - | - | - | - | 100 | - |
| *BZ1-3* | 100 | - | - | - | - | - | - | - | - |
| *Caff. CoA-1\** | 100 | 79 | - | - | 98 | - | - | - | 100 |
| *Caff. CoA-2* | 100 | - | - | - | - | - | - | - | - |
| *Caff. CoA-3* | 100 | 100 | 99 | 98 | 85 | - | - | - | - |
| *CHS* | 100 | - | - | - | 70 | - | - | - | - |
| *DFR-1* | 100 | - | 91 | 100 | - | 92 | - | - | - |
| *DFR-2\*\** | 99 | - | - | - | - | - | - | 99 | - |
| *F3'5'H* | 100 | - | 98 | 86 | 93 | - | - | - | - |
| *F3'H* | 100 | 100 | 93 | 99 | - | - | 83 | - | - |
| Regulatory genes |  |  |  |  |  |  |  |  |  |
| *AN11a* | 100 | - | - | - | - | - | - | - | - |
| *bHLH1* | 100 | 78 | - | 91 | - | - | - | - | - |
| *bHLH2-3* | 100 | - | - | 99 | - | - | - | - | - |
| *bHLH2-2* | 100 | - | 99 | - | - | - | - | - | - |
| *bHLH12* | 100 | 70 | - | 100 | 94 | - | - | - | - |
| *COP1* | 100 | - | - | 99 | - | - | - | - | - |
| *MYB4-1* | 99 | - | - | - | - | - | - | - | - |
| *MYB4-2* | 100 | - | - | - | - | - | - | - | - |
| *MYB16-1* | 100 |  | - | 87 | - | - | - | - | - |
| *MYB16-2* | 100 | - | - | 98 | - | - | - | - | - |
| *SPA1* | 100 | - | - | - | - | - | - | - | - |
| *TTG1* | 100 | - | - | 100 | - | - | - | - | - |

La (*L. arvensis*), Lm (*L. monelli*), O (orange), B (blue)
\* Genes only present in blue petals
\*\* Genes only present in orange petals

**Appendix 1—table 4.** Fixed non-synonymous SNPs in ABP loci that differentiate color morphs and those that differentiate species.

| Gene | Position (aa) | Blue AA | Orange AA | BLOSUM62 |
| --- | --- | --- | --- | --- |
| Fixed Between Color Morphs |  |  |  |  |
| DFR-1 | 279/287<sup>a</sup> | E | Q | +2 |
| MYB4-2 | 361 | M | I | +1 |
| MYB4-2 | 376 | P | S | -1 |
| Fixed Between Species |  |  |  |  |
| BZ1-3 | 435 | K/A<sup>b</sup> | T | -1/0 |
| CaffCoA-3 | 36 | V | A | 0 |
| CaffCoA-3 | 200 | F | Y | +3 |
| CaffCoA-1 (only in blue)<sup>c</sup> | 2 | T | A | 0 |
| CaffCoA-1 (only in blue) | 46 | E | A | -1 |
| CaffCoA-1 (only in blue) | 215 | T | A | 0 |
| CHS | 363 | R | L | -2 |
| F3'H-1 | 19 | S | Y | -2 |
| F3'H-1 | 262 | I | M | +1 |
| F3'H-1 | 324 | Q | H | 0 |
| F3'H-1 | 326 | K | E | +1 |
| F3'H-1 | 338 | A | V | 0 |
| F3'H-1 | 341 | R | L/P | -2/-2 |
| F3'5'H | 2 | E | D | +2 |
| MYB4-1 | 71 | S | T | -1 |
| MYB16-1 | 18 | T | I | +1 |

<sup>a</sup> Two positions separated by a slash indicates the alignment position in *L. arvensis*/*L. monelli* (because of indels).
<sup>b</sup> Two amino acids separated by a slash indicates variable in one species (BLOSUM62 values are reported with a slash in the same order).
<sup>c</sup> “Only in blue” is reported when the locus was not detected in orange samples (still fixed between species, but only for blue).

**Appendix 1—table 5.** Results of hand-crosses between L. arvensis and L. monelli coming from different populations.

| Mother species | Father species | Mother plant* | Father plant* | Mother color | Father color | Fruit production |
| --- | --- | --- | --- | --- | --- | --- |
| *L. monelli* | *L. arvensis* | 630/14_1 | HIN_1.15 | Blue | Blue | 0 |
| *L. monelli* | *L. arvensis* | 630/14_2 | HIN_1.15 | Blue | Blue | 0 |
| *L. monelli* | *L. arvensis* | 630/14_3 | HIN_1.15 | Blue | Blue | 0 |
| *L. monelli* | *L. arvensis* | 630/14_4 | HIN_2.21 | Blue | Blue | 0 |
| *L. monelli* | *L. arvensis* | 630/14_5 | HIN_2.21 | Blue | Blue | 0 |
| *L. monelli* | *L. arvensis* | 630/14_1 | COR_1.6 | Blue | Orange | 0 |
| *L. monelli* | *L. arvensis* | 630/14_2 | COR_1.6 | Blue | Orange | 0 |
| *L. monelli* | *L. arvensis* | 630/14_3 | COR_1.6 | Blue | Orange | 0 |
| *L. monelli* | *L. arvensis* | 630/14_4 | COR_2.5 | Blue | Orange | 0 |
| *L. monelli* | *L. arvensis* | 630/14_5 | COR_2.5 | Blue | Orange | 0 |
| *L. monelli* | *L. arvensis* | 151/14_1 | HIN_1.15 | Blue | Blue | 0 |
| *L. monelli* | *L. arvensis* | 151/14_3 | HIN_1.15 | Blue | Blue | 0 |
| *L. monelli* | *L. arvensis* | 151/14_4 | HIN_2.21 | Blue | Blue | 0 |
| *L. monelli* | *L. arvensis* | 151/14_6 | HIN_2.21 | Blue | Blue | 0 |
| *L. monelli* | *L. arvensis* | 151/14_7 | HIN_8.7 | Blue | Blue | 0 |
| *L. monelli* | *L. arvensis* | 151/14_8 | HIN_8.7 | Blue | Blue | 0 |
| *L. monelli* | *L. arvensis* | 151/14_1 | COR_1.6 | Blue | Orange | 0 |
| *L. monelli* | *L. arvensis* | 151/14_3 | COR_1.6 | Blue | Orange | 0 |
| *L. monelli* | *L. arvensis* | 151/14_4 | COR_1.6 | Blue | Orange | 0 |
| *L. monelli* | *L. arvensis* | 151/14_6 | COR_1.52 | Blue | Orange | 0 |
| *L. monelli* | *L. arvensis* | 151/14_7 | COR_1.52 | Blue | Orange | 0 |
| *L. monelli* | *L. arvensis* | 151/14_8 | COR_1.60 | Blue | Orange | 0 |
| *L. arvensis* | *L. monelli* | HIN_1.15 | 630/14_1 | Blue | Blue | 0 |
| *L. arvensis* | *L. monelli* | HIN_2.21 | 630/14_4 | Blue | Blue | 0 |
| *L. arvensis* | *L. monelli* | HIN_2.21 | 630/14_5 | Blue | Blue | 0 |
| *L. arvensis* | *L. monelli* | COR_1.6 | 630/14_1 | Orange | Blue | 0 |
| *L. arvensis* | *L. monelli* | COR_2.5 | 630/14_5 | Orange | Blue | 0 |
| *L. arvensis* | *L. monelli* | HIN_1.15 | 151/14_1 | Blue | Blue | 0 |
| *L. arvensis* | *L. monelli* | HIN_1.15 | 151/14_3 | Blue | Blue | 0 |
| *L. arvensis* | *L. monelli* | HIN_2.21 | 151/14_4 | Blue | Blue | 0 |
| *L. arvensis* | *L. monelli* | HIN_8.7 | 151/14_8 | Blue | Blue | 0 |
| *L. arvensis* | *L. monelli* | COR_1.6 | 151/14_1 | Orange | Blue | 0 |
| *L. arvensis* | *L. monelli* | COR_1.6 | 151/14_3 | Orange | Blue | 0 |
| *L. arvensis* | *L. monelli* | COR_1.52 | 151/14_6 | Orange | Blue | 0 |
| *L. arvensis* | *L. monelli* | COR_1.60 | 151/14_8 | Orange | Blue | 0 |
| *L. monelli* | *L. arvensis* | 485/14_1 | HIN_1.15 | Orange | Blue | 0 |
| *L. monelli* | *L. arvensis* | 485/14_1 | HIN_1.15 | Orange | Blue | 0 |
| *L. monelli* | *L. arvensis* | 485/14_2 | HIN_2.21 | Orange | Blue | 0 |
| *L. monelli* | *L. arvensis* | 485/14_2 | HIN_2.21 | Orange | Blue | 0 |
| *L. monelli* | *L. arvensis* | 485/14_3 | HIN_8.7 | Orange | Blue | 0 |
| *L. monelli* | *L. arvensis* | 485/14_3 | HIN_8.7 | Orange | Blue | 0 |
| *L. monelli* | *L. arvensis* | 485/14_1 | COR_1.6 | Orange | Orange | 0 |
| *L. monelli* | *L. arvensis* | 485/14_1 | COR_1.6 | Orange | Orange | 0 |
| *L. monelli* | *L. arvensis* | 485/14_2 | COR_1.52 | Orange | Orange | 0 |
| *L. monelli* | *L. arvensis* | 485/14_2 | COR_1.52 | Orange | Orange | 0 |
| *L. monelli* | *L. arvensis* | 485/14_3 | COR_1.60 | Orange | Orange | 0 |
| *L. monelli* | *L. arvensis* | 485/14_3 | COR_1.60 | Orange | Orange | 0 |
| *L. arvensis* | *L. monelli* | HIN_1.15 | 485/14_1 | Blue | Orange | 0 |
| *L. arvensis* | *L. monelli* | HIN_2.21 | 485/14_2 | Blue | Orange | 0 |
| *L. arvensis* | *L. monelli* | HIN_8.7 | 485/14_3 | Blue | Orange | 0 |
| *L. arvensis* | *L. monelli* | COR_1.6 | 485/14_1 | Orange | Orange | 0 |
| *L. arvensis* | *L. monelli* | COR_1.52 | 485/14_2 | Orange | Orange | 0 |
| *L. arvensis* | *L. monelli* | COR_1.60 | 485/14_3 | Orange | Orange | 0 |

\*Population information:
- HIN: *L. arvensis*. Spain, Huelva, Hinojos (37°17'42.6"N, 6°58'05"W) - COR: *L. arvensis*. France, Corsica, Solenazara (41°51'19.6"N, 9°21'43.7"E) - 630/14: *L. monelli*. Spain. Cádiz. Barbate. Breña de Barbate (36°10'39.3"N, 5°58'56.1"W) - 485/14: *L. monelli*. Spain. Tarragona. Mont-Roig (41°5'18"N, 0°56'5"E) - 151/14: *L. monelli*. Portugal. Natural Reserve das Lagoa de St André (38°0'55.2"N, 8°49'11.1"W)

**Appendix 1—table 6.** Climatic differences among morphs. Comparisons indicate significance of differences among morphs (e.g. between the two orange morphs or between the two color morphs within *L. arvensis*) and were Bonferroni adjusted for 76 comparisons (nineteen variables x four morph comparisons).

|  | Variable means (SE) |  |  |  | Morph Comparisons (Kruskal Wallis p values) |  |  |  |
| --- | --- | --- | --- | --- | --- | --- | --- | --- |
|  | arvensis orange | arvensis blue | monelli orange | monelli blue | arvensis difference | orange difference | blue difference | monelli difference |
| AnMnTmp | 11.839 (0.067) | 15.912 (0.093) | 14.508 (0.35) | 15.21 (0.076) | < 0.001 | < 0.001 | < 0.001 | 1.0 |
| MnTmpWrmQtr | 18.71 (0.07) | 22.96 (0.083) | 22.027 (0.32) | 21.659 (0.077) | < 0.001 | < 0.001 | < 0.001 | 1.0 |
| MnTmpCldQtr | 5.534 (0.084) | 9.604 (0.129) | 7.8 (0.432) | 9.464 (0.11) | < 0.001 | < 0.001 | 1.0 | 0.04 |
| AnnPrcp | 768.135 (5.65) | 541.901 (8.085) | 534.675 (25.145) | 663.231 (7.37) | < 0.001 | < 0.001 | < 0.001 | 0.002 |
| PrcpWetMo | 96.007 (0.794) | 87.062 (1.458) | 77.805 (3.642) | 101.085 (1.169) | < 0.001 | 0.003 | < 0.001 | < 0.001 |
| PrcpDryMo | 34.857 (0.427) | 8.609 (0.441) | 8.078 (1.151) | 7.946 (0.243) | < 0.001 | < 0.001 | 1.0 | 1.0 |
| PrcpSeas | 32.321 (0.425) | 59.064 (0.891) | 52.067 (1.748) | 56.902 (0.445) | < 0.001 | < 0.001 | 1.0 | 1.0 |
| PrcpWetQt | 266.99 (2.216) | 234.572 (4.038) | 211.078 (10.287) | 281.789 (3.318) | < 0.001 | < 0.001 | < 0.001 | < 0.001 |
| PrcpDryQt | 121.847 (1.295) | 41.244 (1.606) | 47.494 (4.703) | 42.091 (0.853) | < 0.001 | < 0.001 | 1.0 | 1.0 |
| PrcpWmrQt | 152.319 (1.603) | 51.462 (1.897) | 58.494 (5.645) | 47.816 (0.897) | < 0.001 | < 0.001 | 1.0 | 1.0 |
| PrcpCldQt | 221.595 (2.408) | 206.694 (4.206) | 172.61 (10.381) | 265.361 (3.44) | 0.582 | 0.022 | < 0.001 | < 0.001 |
| DiurnRng | 8.312 (0.038) | 10.038 (0.077) | 10.438 (0.217) | 9.737 (0.08) | < 0.001 | < 0.001 | 1.0 | 0.698 |
| Isoth | 36.593 (0.107) | 40.213 (0.167) | 39.281 (0.491) | 41.305 (0.108) | < 0.001 | < 0.001 | 0.002 | 0.05 |
| TmpSeas | 541.797 (2.946) | 549.092 (4.9) | 584.242 (13.062) | 498.631 (5.108) | 1.0 | 1.0 | < 0.001 | < 0.001 |
| MxTmpWmMo | 24.626 (0.087) | 29.884 (0.11) | 29.34 (0.366) | 28.619 (0.13) | < 0.001 | < 0.001 | < 0.001 | 1.0 |
| MnTmpCoMo | 1.703 (0.081) | 4.712 (0.142) | 2.594 (0.454) | 4.862 (0.126) | < 0.001 | 1.0 | 1.0 | < 0.001 |
| TmpAnRng | 22.923 (0.104) | 25.172 (0.195) | 26.747 (0.56) | 23.757 (0.219) | < 0.001 | < 0.001 | < 0.001 | < 0.001 |
| MnTmpWetQtr | 10.711 (0.098) | 12.542 (0.128) | 11.798 (0.465) | 10.655 (0.097) | < 0.001 | 1.0 | < 0.001 | 1.0 |
| mnTmpDryQtr | 13.224 (0.178) | 21.795 (0.179) | 21.802 (0.297) | 21.399 (0.08) | < 0.001 | < 0.001 | 1.0 | 1.0 |

**Appendix 1—table 7.** Logistic regression models for each species on a subset on non-correlated climate variables. The best models using AIC for *L. arvensis* (Arv) and *L. monelli* (Mon) are presented. “Good models” have lower, but close AIC values and reinforce the importance of the variables included in the best model.

| Coefficients | Estimate | Std. Error | z value | p |
| --- | --- | --- | --- | --- |
| Intercept | 14.5510859 | 0.8911108 | 16.33 | < 0.0001 |
| PrcpSeas | -0.0453151 | 0.0028234 | -16.05 | < 0.0001 |
| Isoth | -0.1980946 | 0.0168407 | -11.76 | < 0.0001 |
| TmpSeas | -0.0070202 | 0.0005796 | -12.11 | < 0.0001 |

***L. arvensis* Good Models**
| Intercept | Isoth | MeanTemp<br>WarmQtr | PrcpSeas | PrcWarm<br>Mon | TmpSeas | df | Delta<br>AIC |
| --- | --- | --- | --- | --- | --- | --- | --- |
| 47.92 | 20.240 |  | -37.950 | 2.044 | -46.670 | 5 | 1.20 |
| 58.550 | 18.270 | 1.649 | -39.18 |  | -57.010 | 5 | 1.96 |
| 45.220 | 19.610 | 5.531 | -37.94 | 2.695 | -44.090 | 6 | 2.80 |

***L. monelli* Best Model**
| Coefficients | Estimate | Std. Error | z value | p |
| --- | --- | --- | --- | --- |
| Intercept | -1.568856 | 0.635342 | -2.469 | 0.0135 |
| PrcpWetMo | -0.035533 | 0.005195 | -6.840 | < 0.0001 |
| MnTmpWetQtr | 0.229809 | 0.049652 | 4.628 | < 0.0001 |

***L. monelli* Good Models**
| Intercept | Isoth | MeanTemp<br>WarmQtr | PrcpSeas | PrcWarm<br>Mon | TmpSeas | df | Delta<br>AIC |
| --- | --- | --- | --- | --- | --- | --- | --- |
| 22.280 | -81.070 | 215.70 | 20.42 | -24.93 | -20.80 | 6 | 1.68 |
| 46.480 | -89.320 | 210.80 |  | -24.22 | -44.92 | 5 | 5.67 |
| -0.02067 | -70.320 | 210.30 |  | -17.75 |  | 4 | 5.82 |

**Appendix 1—table 8.** Significance values from randomization tests indicating whether orange or blue populations are more different from one another climatically than expected by chance.

|  | Orange<br>divergent | Blue divergent | Conclusion |
| --- | --- | --- | --- |
| AnMnTmp | < 0.001 | 0.999 | Orange more divergent |
| MnTmpWrmQtr | < 0.001 | 0.999 | Orange more divergent |
| MnTmpCldQtr | < 0.001 | 0.999 | Orange more divergent |
| AnnPrcp | < 0.001 | 0.999 | Orange more divergent |
| PrcpWetMo | 0.205 | 0.794 | Not Significant |
| PrcpDryMo | < 0.001 | 0.999 | Orange more divergent |
| PrcpSeas | < 0.001 | 0.999 | Orange more divergent |
| PrcpWetQt | 0.039 | 0.96 | Orange more divergent |
| PrcpDryQt | < 0.001 | 0.999 | Orange more divergent |
| PrcpWmrQt | < 0.001 | 0.999 | Orange more divergent |
| PrcpCldQt | 0.796 | 0.203 | Not significant |
| DiurnRng | < 0.001 | 0.999 | Orange more divergent |
| Isoth | < 0.001 | 0.999 | Orange more divergent |
| TmpSeas | 0.702 | 0.297 | Not significant |
| MxTmpWmMo | < 0.001 | 0.999 | Orange more divergent |
| MnTmpCoMo | 0.410 | 0.958 | Not significant |
| TmpAnRng | < 0.001 | 0.999 | Orange more divergent |
| MnTmpWetQtr | 0.999 | < 0.001 | Blue more divergent |
| MnTmpDryQtr | < 0.001 | 0.999 | Orange more divergent |

**Appendix 1—table 9.**
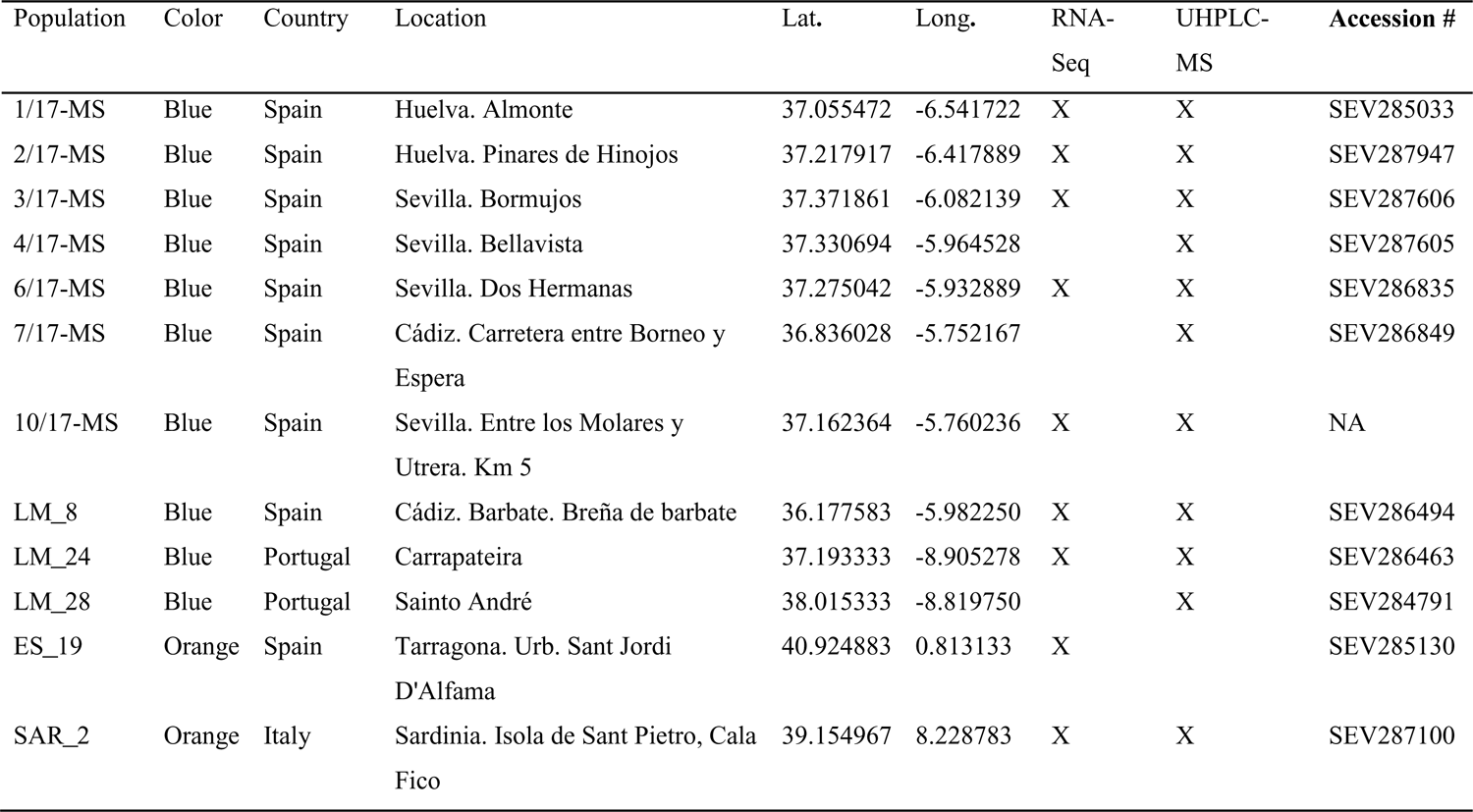
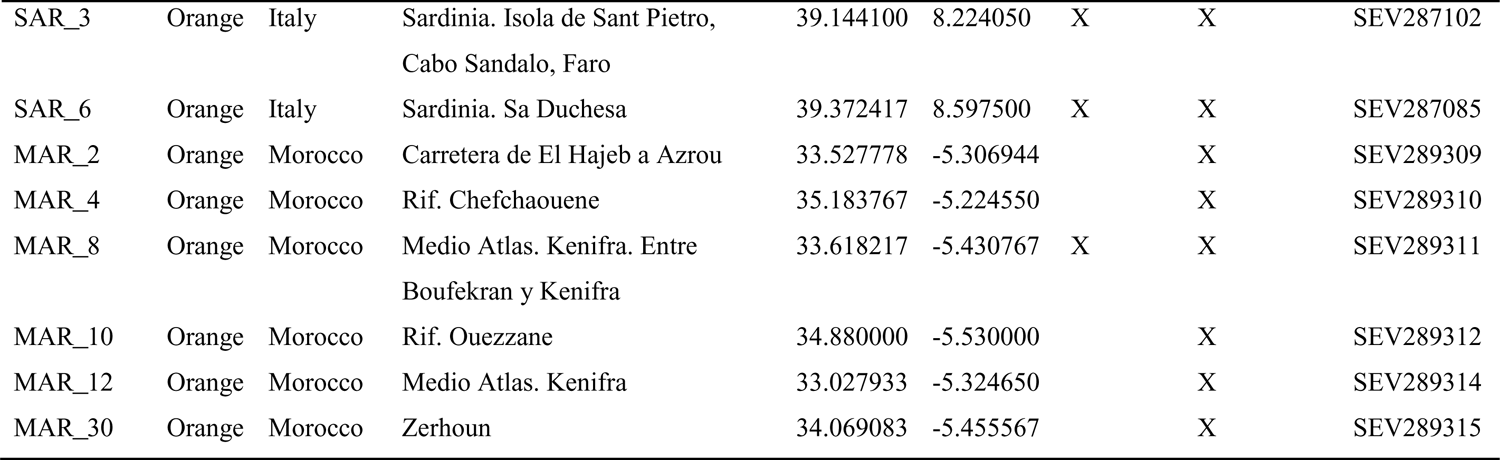
Plant population locations and herbarium accession numbers for *Lysimachia monelli* samples used for RNA-seq and UHPLC-MS (flavonoid profiling) (marked with a X). Voucher specimens were deposited in the University of Seville Herbarium (SEV).

**Appendix 1—table 10.** Summary of sequencing and assembly results for the 20 *L. monelli* petal samples.

| Sample | Population | Flower color | Number of reads | Number of filtered reads | Filtered reads aligned to the assembly (%) | Number of transcripts | Number of trinity genes* |
| --- | --- | --- | --- | --- | --- | --- | --- |
| Lymo_1 | ES_19 | orange | 19,356,011 | 18,421,349 | 92.48 | 47,377 | 26,986 |
| Lymo_2 | SAR_3 | orange | 17,662,750 | 16,866,036 | 93.30 | 45,323 | 27,137 |
| Lymo_3 | SAR_2 | orange | 18,038,810 | 17,237,553 | 93.16 | 48,194 | 29,126 |
| Lymo_7 | SAR_6 | orange | 17,084,149 | 16,312,231 | 92.11 | 49,194 | 29,265 |
| Lymo_18 | SAR_6 | orange | 17,166,042 | 16,255,988 | 90.86 | 48,459 | 28,927 |
| Lymo_23 | MAR_8 | orange | 16,537,229 | 15,503,176 | 92.47 | 48,701 | 28,213 |
| Lymo_24 | MAR_8 | orange | 16,709,072 | 15,886,953 | 92.23 | 49,265 | 28,074 |
| Lymo_34 | ES_19 | orange | 17,230,125 | 16,466,487 | 92.21 | 49,477 | 28,830 |
| Lymo_35 | ES_19 | orange | 23,467,043 | 22,697,727 | 92.73 | 46,959 | 27,097 |
| Lymo_36 | SAR_2 | orange | 16,981,334 | 15,773,601 | 92.57 | 52,169 | 31,542 |
| Lymo_40 | 2/17-MS | blue | 18,359,772 | 17,602,178 | 93.03 | 46,380 | 26,228 |
| Lymo_45 | LM_8 | blue | 17,828,507 | 16,984,718 | 92.32 | 48,951 | 28,143 |
| Lymo_8 | LM_24 | blue | 16,850,071 | 16,124,259 | 91.26 | 45,973 | 26,624 |
| Lymo_9 | 1/17-MS | blue | 18,050,225 | 17,338,160 | 92.84 | 45,891 | 26,354 |
| Lymo_11 | 6/17-MS | blue | 19,073,958 | 18,182,529 | 92.42 | 47,224 | 26,815 |
| Lymo_12 | LM_8 | blue | 18,350,299 | 17,810,293 | 93.54 | 44,695 | 25,856 |
| Lymo_13 | 10/17-MS | blue | 17,255,892 | 16,498,600 | 92.63 | 46,720 | 26,591 |
| Lymo_17 | 3/17-MS | blue | 17,405,546 | 16,803,173 | 93.02 | 47,313 | 27,096 |
| Lymo_25 | 3/17-MS | blue | 16,982,762 | 15,953,718 | 92.29 | 48,271 | 27,650 |
| Lymo_29 | 6/17-MS | blue | 16,331,233 | 15,578,920 | 93.01 | 45,869 | 25,935 |

\* Trinity genes were limited to those having TMM > 1.

